# Genetic Modulation of Protein Expression in Rat Brain

**DOI:** 10.1101/2024.02.17.580840

**Authors:** Ling Li, Zhiping Wu, Andrea Guarracino, Flavia Villani, Deihui Kong, Ariana Mancieri, Aijun Zhang, Laura Saba, Hao Chen, Hana Brozka, Karel Vales, Anna N. Senko, Gerd Kempermann, Ales Stuchlik, Michal Pravenec, Pjotr Prins, Junmin Peng, Robert W. Williams, Xusheng Wang

## Abstract

Genetic variations in protein expression are implicated in a broad spectrum of common diseases and complex traits. However, the fundamental genetic architecture and variation of protein expression have received comparatively less attention than either mRNA or classical phenotypes. In this study, we systematically quantified proteins in the brains of a large family of rats using tandem mass tag (TMT)-based quantitative mass-spectrometry (MS) technology. We identified and quantified a comprehensive proteome of 8,119 proteins from Spontaneously Hypertensive (SHR/Olalpcv), Brown Norway with polydactyly-luxate (BN-Lx/Cub), and 29 of their fully inbred HXB/BXH progeny. Differential expression (DE) analysis identified 597 proteins with significant differences in expression between the parental strains (fold change > 2 and FDR < 0.01). We characterized 95 variant peptides by proteogenomics approach and discovered 464 proteins linked to strong *cis*-acting quantitative trait loci (pQTLs, FDR < 0.05). We also explored the linkage of pQTLs with behavioral phenotypes in rats and examined the sex-specific pQTLs to reveal both distinct and shared *cis*-pQTLs between sexes. Furthermore, by creating a novel view of the rat pangenome, we improved the ability to pinpoint candidate genes underlying pQTL. Finally, we explored the connection between the pQTLs in rat and human disorders, underscoring the translational potential of our findings. Collectively, this work demonstrates the value of large and systematic proteo-genetic datasets in understanding protein modulation in the brain and its functional linkage to complex central nervous system (CNS) traits.

## Introduction

Genetic linkage analyses in rodents have unveiled numerous loci contributing to a diverse array of traits, including susceptibility to addictive behaviors, neurodegeneration, and hypertension ^1–3^. Despite these discoveries, linkage analyses alone fail to dissect the underlying biological processes causing these traits ^3,4^. However, a growing body of converging evidence strongly indicates that the associations between genomic loci and traits are primarily mediated through the genetic modulation of expression on molecular endophenotypes ^5,6^. The endophenotypes encompass a wide spectrum of molecules—mRNAs, proteins, and metabolites—that serve as intermediates in complex cascades that link DNA variants to higher order traits. The accuracy and throughput with which molecular endophenotypes can now be quantified in larger populations and families holds significant promise in bridging the gap between genomic loci and traits ^7,8^.

Almost all work on the genetics of expression and its linkage to higher order traits has been involved in mRNA expression first using arrays and now using RNA sequencing ^9–11^. In contrast, much less is known about the arguably more functionally relevant variation in the genetic control of protein expression and its causal contributions to higher order traits. Variations in mRNA expression often do not reliably reflect variations in protein expression due to intricate post-transcriptional and translational regulatory mechanisms ^12–15^. As the main driver of cellular function ^16^, variation in protein levels should have a more direct impact on cell and tissue function. Consequently, unraveling the genetic variation and the genetic control of protein levels holds great potential to dissect molecular and cellular mechanisms underlying the variation in traits.

To investigate the genetic basis of trait variation in molecular endophenotypes, genetic reference populations, such as the mouse BXDs ^17–19^, Collaborative Cross ^20,21^, and the HXB/BXH recombinant inbred (RI) family ^22,23^, have been used extensively over the past decade, but surprisingly never to study protein levels in the brain in any species other than humans ^24^. The HXB/BXH family is the largest and most deeply phenotyped set of genetically diverse but fully inbred set of rats. It has been used mainly to dissect loci genetic variants that modulate metabolic and cardiovascular diseases ^23,25–27^, but in recent work this family is increasingly being used to study the genetics of substance use disorders ^28–30^. All members of the family have been fully sequenced, and high-density marker maps of all classes of DNA variants are now available for both forward and reverse genetic analyses ^31,32^. There are also extensive transcriptomic data sets for several systems ^33–35^, but now for the first time also for the brain ^28^. Finally, there is extensive and curated phenotypic data for this family in GeneNetwork.org ^36–38^, and this enables on-line mapping, as well as integrative analyses to explore potential networks linking loci to traits.

In this study, we systematically quantify the variation and genetic architecture of protein expression in whole brain and behavioral traits in active place avoidance as a task critically dependent on the hippocampus in the rat HXB/BXH RI strains. We start by generating proteome across 29 HXB/BXH RI strains, in conjunction with the two parental strains, SHR and BN-Lx. We then quantify sequence and expression variations at the protein level between the two parental strains using a proteogenomics approach. We identify both *cis* and *trans*-acting loci contributing to variation in protein expression in the brain. We zoom in on gene candidates using pangenome tools. Finally, we explore relations between gene loci, mRNA and protein expression, and phenotypic traits.

## Results

### Proteomic profiling and analysis of the HXB/BXH RI rat family

In our study, we explored genetic variation in protein expression, identified protein expression quantitative trait loci (pQTLs), uncovered sex-specific pQTLs, assessed the impact of pQTLs on phenotypic traits, revealed causal variants within the rat pangenome, and investigated potential links between rat pQTLs and human disorders (**Figure 1**). For the proteomics experiments, we employed two batches of 11-plex and three batches of 16-plex tandem mass tags (TMT) coupled with two-dimensional liquid chromatography tandem mass spectrometry (LC/LC-MS/MS) strategy (**Figure 2A**). In this analysis, we identified and quantified a total of 11,118 proteins in at least one batch at the protein false discovery rate (FDR) <1%. Out of these, 8,124 (73.07%) proteins were detected across all 62 samples (**Figure S1, Figure S2A** and **Table S1**). We performed extensive quality controls, including sample identity verification using the SMAP program ^16^ and batch effect removal using the LIMMA package ^39^. Principal component analysis (PCA) showed that two replicates of the two parental rat strains grouped well (**Figure 2B**). Pearson correlation analysis showed that the two replicates of each strain displayed a high correlation (*r* > 0.99; **Figure S2B**). Paired *t*-test of 31 male-female littermates revealed that only 60 (0.7%) showed differences between sexes (*p* < 0.01; before adjusting for multiple correction) **(Figure S2C**). In contrast, the two parental strains exhibited a large variation in protein expression (**Figure S2D**).

**Figure 1.**
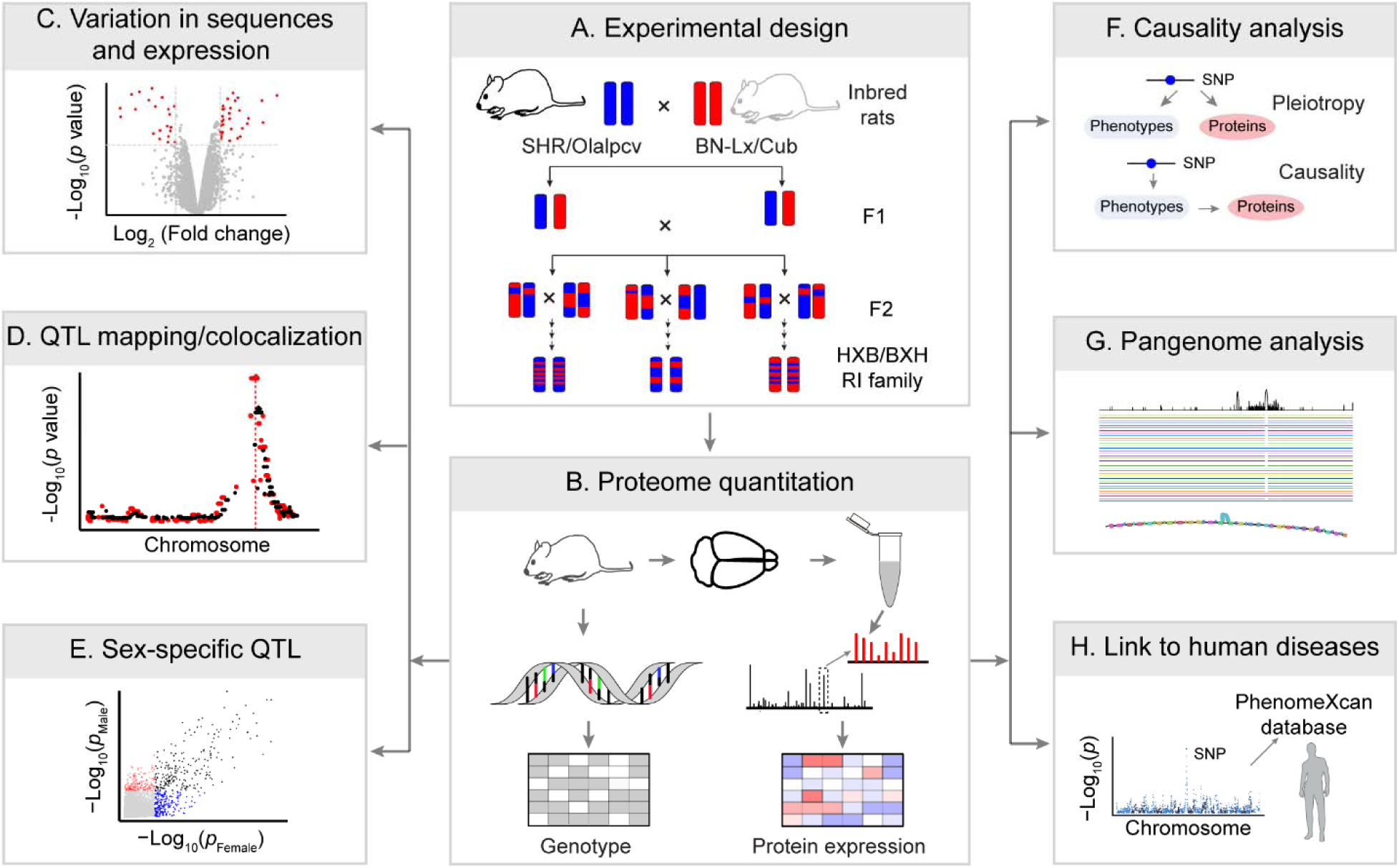
Experimental design and analysis pipeline employed in this study. (A) Schematic representation illustrating the inclusion of postmortem brain samples from 62 participants, encompassing 29 HXB/BXH R1 strains alongside their respective parental strains. (B) Profiling of the deep brain proteome was conducted using 11-plex/16-plex TMT-based proteomics, followed by rigorous quality control and comprehensive data analysis. The brain proteomic data, along with corresponding genotype data, were meticulously prepared for subsequent linkage analysis. (C) Volcano plot demonstrates the differential expression of proteins. (D) QTL analysis was employed to identify genetic regulation of protein expression or gene expression, colocalization *cis*-pQTL and *cis*-eQTL, or *cis*-pQTL and *trans*-pQTL. (E) QTL analysis employed to identify genetic regulations of protein expression in different sex. (F) Integrative analysis to link pGenes to phenotypes. (G) Integrative analysis to link pGenes to phenotypes. (H) Pangenome analysis to explore the variation across all mapping strains.

**Figure 2.**
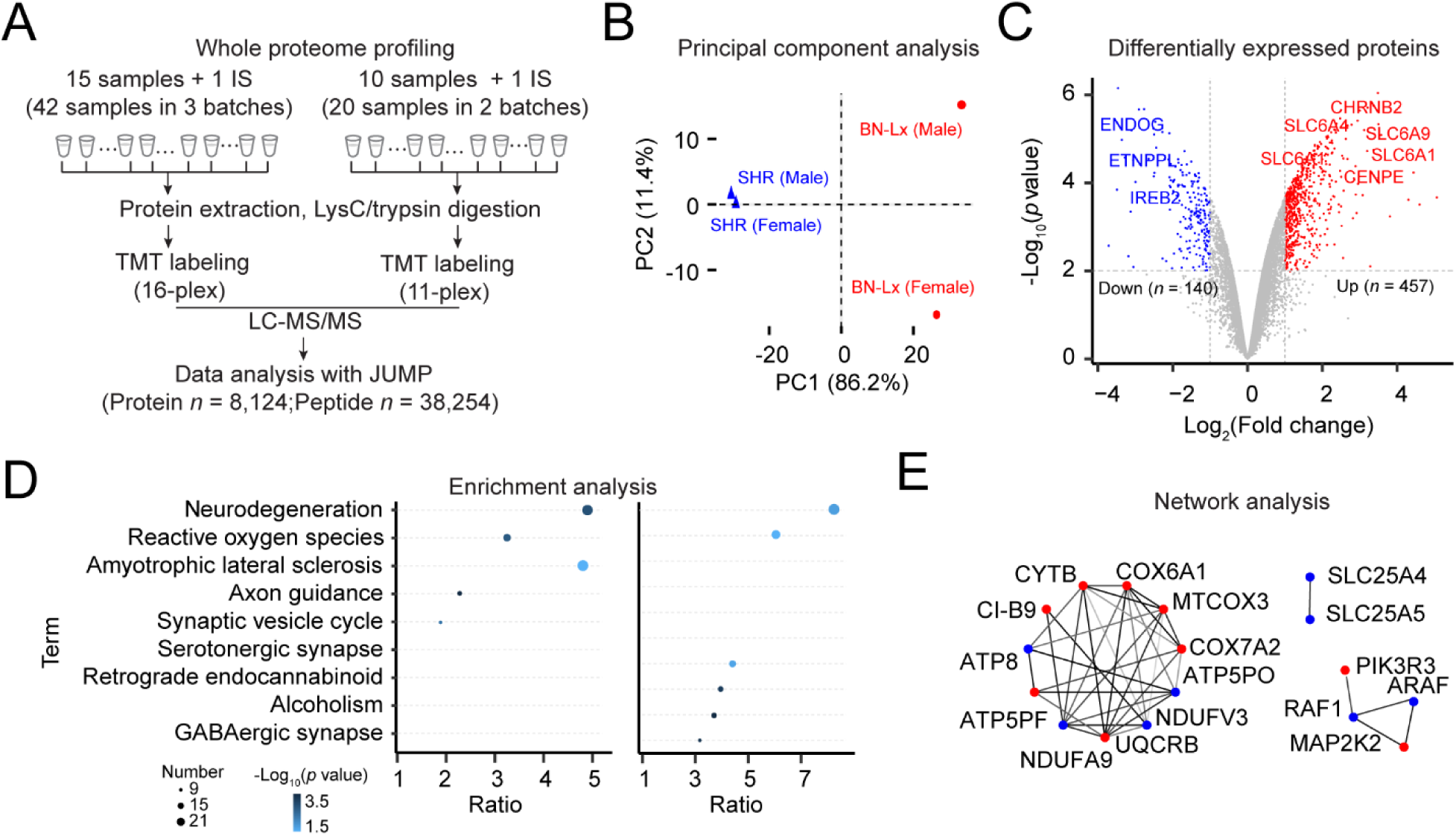
Deep profiling of rat brain proteome. (A) Workflow of 11-plex/16-plex TMT-based proteome analysis. LC/LC-MS/MS was performed on a total of 10/15 samples, along with 1 internal standard (i.e., 10/15 pooled samples). MS raw data were analyzed using JUMP software. (B) Principal-component analysis (PCA) of all quantified proteins. (C) Volcano plot displaying the differentially expressed proteins. Log_2_ fold change was plotted against the −log_10_ adjusted *p*-value with one, criterion: 2-fold change and 1% adjusted *p* values. (D) KEGG pathways enrichment analysis of DE proteins, Higher in SHR (left), Higher in BN-Lx (right). (E) Network of proteins in the ROS (Reactive Oxygen Species) pathway.

As the transcriptomic data had previously been generated for the HXB/BXH RI strains ^40^, we compared our proteomic data to transcriptomic findings. The quantified proteins cover ∼70% of the range of transcripts detected by the transcriptomic data (**Figure S3A**). A moderate positive correlation between expression levels of mRNAs and proteins (*r* = 0.35) was observed (**Figure S3B**), which is consistent with previous findings in human study ^24^.

### Genetic variation in sequences and expression levels

Our deep proteome provides an opportunity to investigate sequence variations between the two parental strains at the protein level. To identify variant peptides with the SHR allele, we performed proteogenomic analysis using JUMPg ^41^. We identified a total of 95 variant peptides at a peptide FDR of 1% (**Table S2A**). Of these variant peptides, we also detected 61 corresponding reference peptides with the BN-Lx reference allele. As expected, all variant peptides showed significantly higher expression in SHR compared to BN-Lx, as illustrated with two variant peptides (**Figure S4)**.

To measure variations in protein expression between the two parental strains of the HXB/BXHRI family—SHR and BN-Lx strains—we conducted a differential protein expression analysis using the LIMMA package. We identified a total of 597 differentially expressed proteins (DEPs) with an adjusted *p* value of 0.01 and a log_2_ fold change (log_2_FC) of 1 or greater (**Figure 2C** and **Table S2B**). Among these, 457 proteins displayed higher expression in the SHR strain, while 140 proteins exhibited higher expression in the reference BN-Lx strain. Kyoto Encyclopedia of Genes and Genomes (KEGG) enrichment analysis revealed that DEPs with higher expression in SHR were significantly enriched in terms related to reactive oxygen species (ROS), synaptic vesicle cycle, and amyotrophic lateral sclerosis (ALS) (**Figure 2D**, **Tables S2C**, and **Table S2D**). By projecting these DEPs to the STRING protein-protein interaction (PPI) network (version 12), three functional modules of the DEPs in the ROS pathway were identified (**Figure 2E**). The DEPs with higher expression in BN-Lx showed significant enrichment in the metabolic pathway.

A subset of proteins—391 out of 8,268—exhibited remarkably high variability, operationally defined as two standard deviations above the average coefficient of variation (CV, mean = 1.2 x 10^−2^; SD = 8.6 x 10^−3^) (**Figure S5A** and **Table S1**). These highly variable proteins were significantly enriched in pathways associated with Serotonergic synapse (including CYP2D10, GNG4, CASP3, CYP2D3, GNG8, CYP2D1, PLCB2, CYP2D26), oxidative phosphorylation (including NDUFA8, NDUFAB1, COX17, ATP5PO, CYCT, ATP6V0C, ATP5F1E, COX6B1), and inflammatory mediator regulation of TRP channels (such as PRKCH, MAP1, ASIC2, PLCB2, KNG1), (**Figure S5B** and **Table S2E**) ^42–45^.

We next sought to estimate the heritability (*h*^2^) of protein expression, which is defined as the proportion of additive genetic variation contributing to the overall observed variation. The analysis revealed that the protein heritability in the HXB/BXH family has a median heritability of 71%, ranging from 0.08 to 0.99 across all expressed proteins (**Figure 3A** and **Table S1**). For example, 2,4-dienoyl-CoA reductase 1 (DECR1) showed a high heritability of 0.94. The high heritability of proteins in the HXB/BXH family makes them particularly amenable to pQTL mapping.

**Figure 3.**
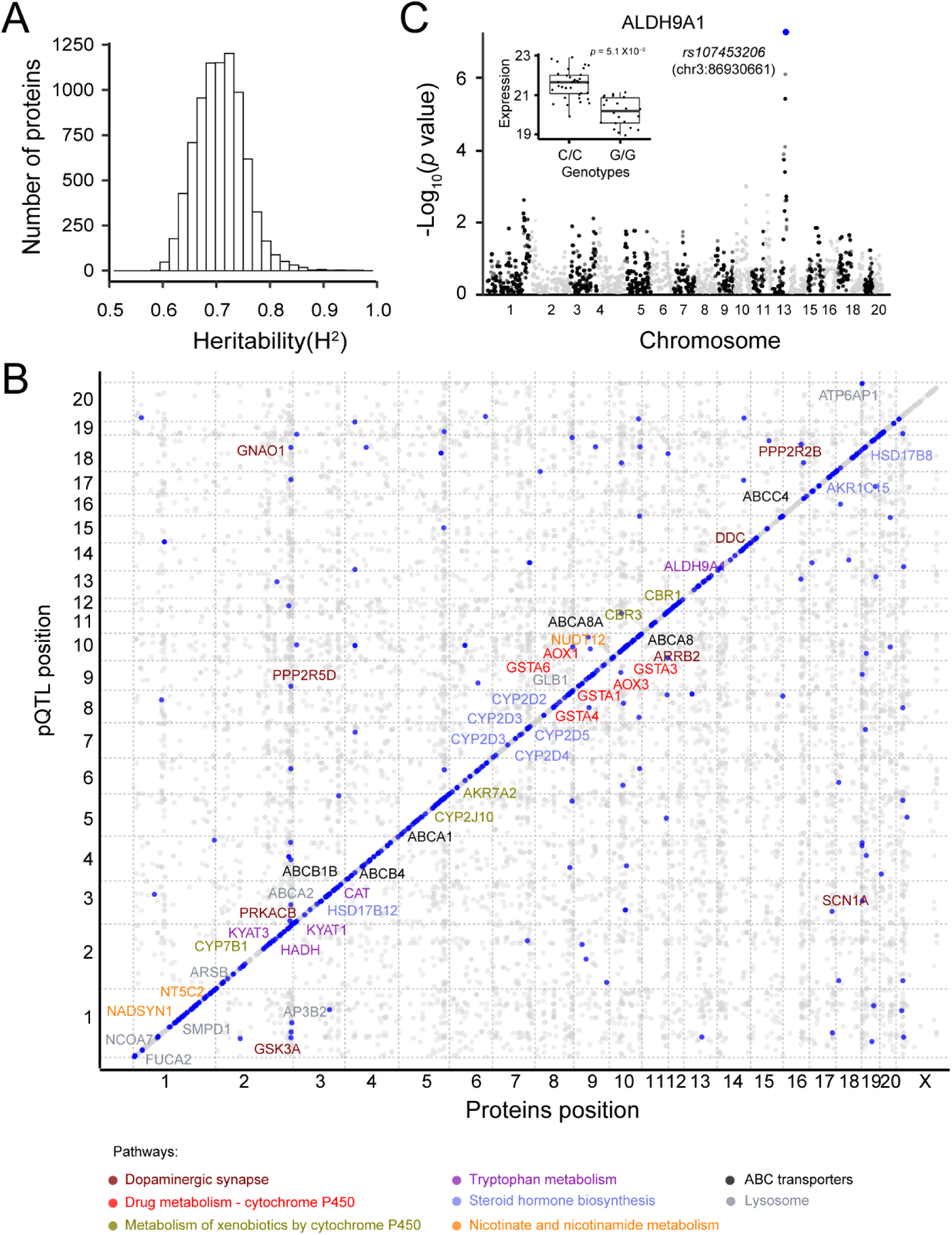
Genetic modulation of the brain proteome. (A) Genetic variance and heritability estimate for all 8,119 identified proteins. (B) Scatter plot displaying the physical gene location in total megabases versus QTL genetic location in total centimorgans. (C) Manhattan plot showing a *cis*-pQTL (i.e., *rs107453206*) is associated with ALDH9A1 protein expression.

### Genetic modulation of protein expression

We proceeded to investigate the genetic modulation of protein expression by conducting a comprehensive proteome-wide linkage analysis. Leveraging data from 8,119 proteins detected in 29 HXB/BXH RI strains and 10,465 genotypes, we identified 318 *cis*-acting (or local acting) loci that modulate the expression of 464 cognate proteins at a cutoff value *p* < 1 × 10^−3^, (see methods, **Figure 3B, Figure S6** and **Table S3A**) using the GEMMA mapping software. These *cis*-pQTLs were enriched for metabolic pathways, such as drug and amino acid metabolisms. For example, *cis*-pGenes such as AOX3, CAT, KYAT3, AOX1, KYAT1, HADH, ALDH9A1 were enriched in the Tryptophan metabolism (**Figure 3B**). As an example, we found that ALDH9A1 (UniProt ID: Q9JLJ3) had a *cis*-pQTL with a linkage score *p* value of 4.76 × 10^−8^ (**Figure 3C**), located on Chromosome 13 at 86,930,661 bp (*rs107453206*). Comparing the alternative allele (*C/C*) with the reference allele (*G/G*), we observed a significant increase in protein expression (20.13 *vs* 21.56, *p* = 5.10 × 10^−9^) (**Figure 3C inset**). ALDH9A1 exhibits high expression levels in brain ^46,47^ and plays a role in an alternate biosynthesis pathway of gamma-aminobutyric acid (GABA) by catalyzing the oxidation of γ-aminobutyraldehyde ^47–49^.

To explore whether the linkage between the locus, mRNA levels, and corresponding protein expression, we compared 18,278 genes expression quantitative trait loci (eQTLs) using companion brain RNA-seq data for 30 HXB/BXH RI cases ^40^. We identified 1,247 *cis*-eQTLs (**Table S4**) with acceptance cutoff thresholds (*p* = 3.00 × 10^−3^; see methods). We were able to compare these with 318 *cis*-pQTLs and found 115 pairs of colocalized QTLs (**Figure 4A** and **Table S5**). The vast majority (75.6%; 87/115) of these co-localized mRNA-protein loci showed the same allelic effect polarity, with a mean Pearson correlation coefficient (*r*) of 0.78 (*p* value = 2.2 × 10^−16^; **Figure 4B**). This strongly supports the notion that an increase in the effect size of *cis*-eQTLs is indicative of an increase in that of *cis*-pQTLs. In contrast, we observed only a moderate correlation of expression levels between colocalized proteins and genes (*r* = 0.43; *p* value = 1.23× 10^−6^; **Figure 4C**). For example, AKR1B8 (UniProt ID: Q6AY99) was identified as a significant *cis*-pQTL (*p* value = 2.03 × 10^−10^), and a significant *cis*-eQTL (*p* value = 5.50 × 10^−18^) (**Figure 4D**), supported by a significant positive correlation of expression at the gene and protein levels (**Figure 4E**).

**Figure 4.**
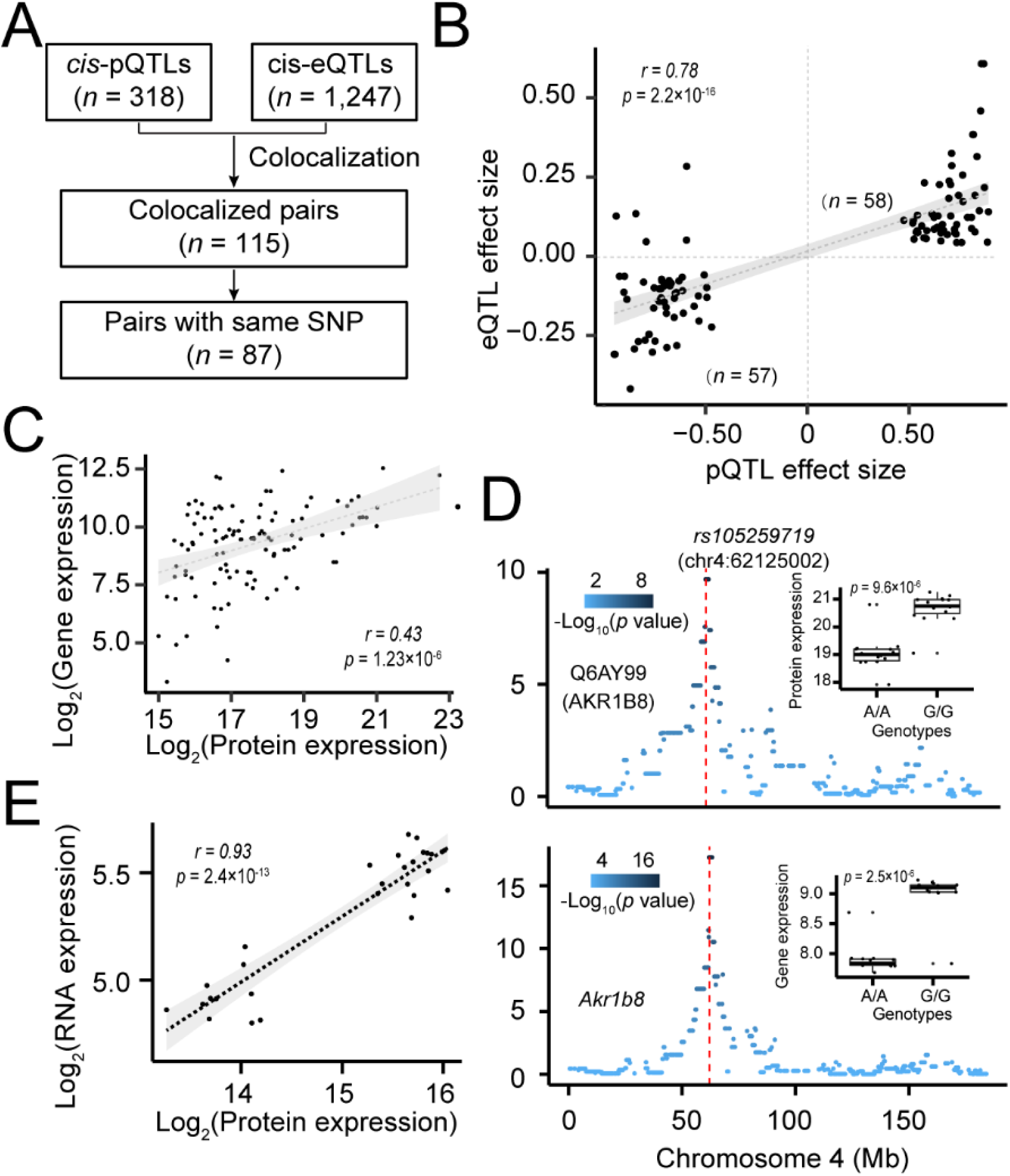
**Co-localization of QTLs regulating the expression level of genes and proteins**. (A) Venn diagram shows number of genes colocalized by *cis*-eQTLs and *cis*-pQTLs. (B) Scatter plot showing the distribution of effect sizes of colocalized *cis*-eQTLs and *cis*-pQTLs. (C) Correlation of expression levels of colocalized *cis*-eGenes and *cis*-pGenes. (D) Scatter plot showing a colocalized QTL modulating *Akr1b8* gene and protein expression. (E) Correlation of expression levels of colocalized *cis*-eGenes and *cis*-pGenes.

### Sex-specific modulation of protein expression

To investigate the impact of sex on protein expression and its genetic regulation, we employed male and female protein expression data to conduct a proteome-wide linkage analysis. Utilizing expression levels of 8,119 proteins from both male and female HXB/BXH RI strains, we identified 305 *cis*-acting loci in females modulating the expression of 433 proteins (**Table S3B**, with a cutoff value of *p* < 3.33 × 10^−3^), and 296 *cis*-acting loci in male influencing the expression of 405 proteins (**Table S3C**), with a cutoff value of *p* < 2.85 × 10^−3^ (**see methods**). Of these, 219 proteins with *cis*-pQTLs overlapped between males and females, while 214 proteins were identified exclusively in females and 186 proteins exclusively in males (**Figure S7A**). Furthermore, when comparing these findings with 318 proteins identified from sex-averaged expression levels, we observed 134 (42%) proteins that were shared by females, males, and sex-average (**Figure S7B**). Enrichment analysis revealed enrichment in multiple sex-related pathways, such as the regulation of peptide hormone secretion and intracellular estrogen receptor signaling pathways. For instance, the expression of PFKL (Phosphofructokinase Liver Type) in the modulation of peptide hormone secretion pathway is controlled by a locus on Chromosome 20 (*rs64542764*, Chr20:10,518,616), with a mapping *p*-value of 3.46 × 10^−4^ (**Figure S7C**). PFKL was previously found to be sex differences in the metabolism of glucose and glycerol ^50^. This analysis highlights the role of the sex-specific genetic factors in shaping protein expression patterns.

### *Trans*-pQTLs modulate expression of downstream proteins

To determine whether a *cis*-pQTL regulates the expression of distant proteins through its colocalized *trans*-pQTL, we conducted a linkage analysis for *trans*-pQTLs (> 10 Mb away from the regulated protein on the same chromosome or different chromosomes), identifying 87 *trans*-acting loci that modulate 116 proteins’ expression (**Table S3A**, permutation *p* value < 10^−5^). By colocalizing these 87 *trans*-pQTLs with 318 *cis*-pQTLs, we found 45 pairs of colocalized QTL signals (**Table S6**). One colocalized QTL is located on Chromosome 3 near the marker *rs107535368* (Chr3:11,913,651 bp). This locus regulates the expression of FLOT1 (flotillin 1) as a *trans*-pQTL, with a mapping *p* value of 1.43 × 10^−7^, and the expression of LRSAM1 (i.e., leucine rich repeat and sterile alpha motif containing 1) as a *cis*-pQTL, with a mapping p value of 1.48 × 10^−10^ (**Figure 5A**). Despite the absence of an apparent functional link between LRSAM1 and FLOT1, LRSAM1 emerges as a plausible candidate protein regulating the protein expression of FLOT1 (**Figure 5B**).

**Figure 5.**
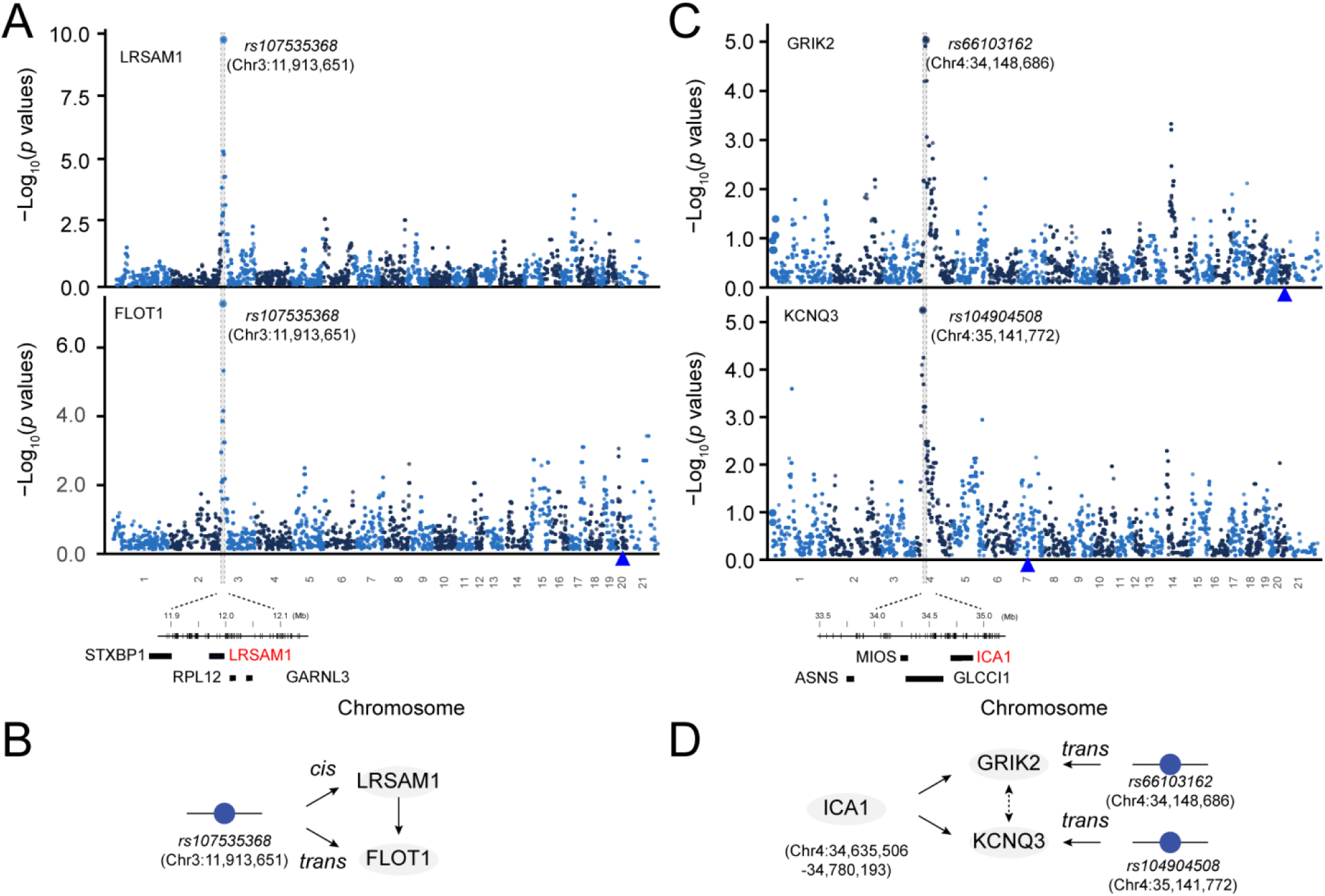
Example of genetic regulation of a protein with *trans*-pQTL. (A) Manhattan plot showing a SNP (*rs107535368*) reveals its association with LRSAM1 protein expression through *cis*-regulation, and simultaneously *trans*-regulates FLOT1 protein expression. The x-axis represents genomic positions, while the y-axis denotes the –log_10_-transformed *p*-values of the association. The QTL region is highlighted in the grey box, covering four candidate genes, including LRSAM1 protein. (B) Schematic diagram showing the regulation of LRSAM1 abundance by a *cis*-pQTL, which regulates FLOT1 protein expression level as a *trans*-QTL. (C) Manhattan plot showing a *trans*-pQTL (i.e., *rs66103162*) associated with GRIK2 protein expression and another *trans*-pQTL (i.e., *rs104904508*) associated with KCNQ3 protein expression. (D) Schematic representation elucidating the regulatory relationships of the two proteins, GRIK2 and KCNQ3, influenced by a *trans*-pQTLs, as well as a candidate gene, ICA1.

Another notable *trans*-pQTL example is a locus on Chromosome 4 regulating the expression of GRIK2, with a mapping *p* value of 8.09 × 10^−6^, and another locus on Chromosome 4 regulating the expression of KCNQ3, with a mapping *p* value of 5.61 × 10^−6^ (**Figure 5C**). By examining the candidate proteins within the locus, we found that ICA1 is a compelling candidate protein regulating the expression of both GRIK2 and KCNQ3. The connection is supported by the protein-protein interaction (PPI) network between the three proteins (**Figure 5D)**.

### Linking variation in protein expression to phenotypic traits

To determine whether a potential causal association between pQTLs, as identified in this study, and phenotypic traits exists, we collected eight behavioral traits for the HXB/BXH family that are associated with spatial learning (**Table S7**). Spatial learning was assessed with an active place avoidance task with reversal (a dry-land spatial task), open-field, and beam-walking tests. Linkage analysis revealed that a locus (i.e., *rs105225151;* Chr10:48,949,496 bp) is significantly associated with Kurtosis factor of annular-Gaussian beam, with a mapping *p* value of 1.86 x 10^-3^ (**Figure 6A, B**). The locus was also found to be associated with the expression level of TRPV2 protein, with a mapping *p* value of 7.09 ×10^−6^ (**Figure 6C**). The reference allele has 1.72-fold (20.24 *vs* 19.46 on log_2_ scale) higher expression than the alternative allele (**Figure 6D**). A Bayesian Network analysis revealed that this locus potentially has a causal effect on both protein expression and phenotype (**Figure 6E**) using the Bayesian Network Webserver (BNW) program^51,52^.

**Figure 6.**
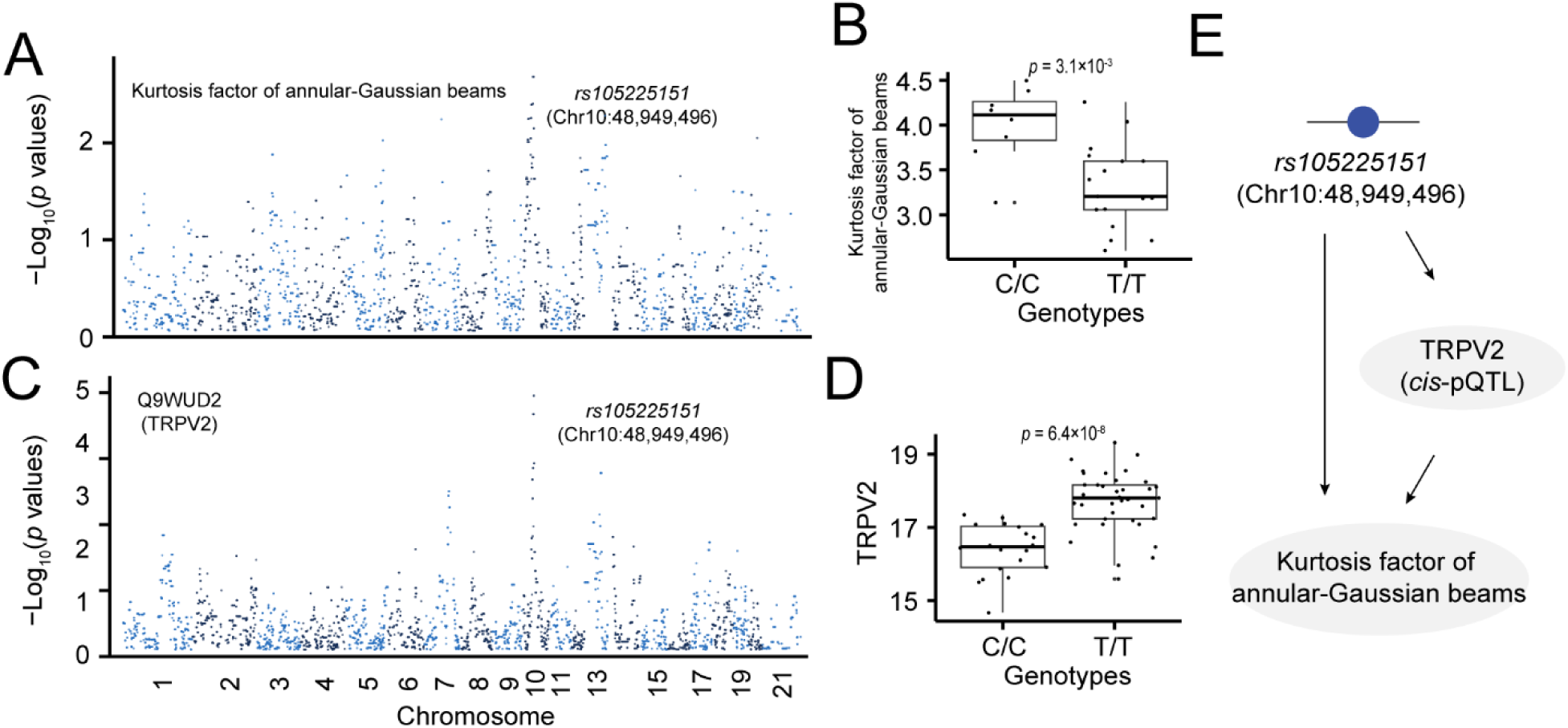
Association of proteins with *cis*-pQTL and behavioral traits. (A) Manhattan plot showing the association of a QTL (i.e., *rs105225151*) with a behavioral trait: Kurtosis factor of annular-Gaussian beams. (B) Box plot showing the trait values between reference allele and the alternative allele. (C) Manhattan plot showing a *cis*-pQTL (i.e., *rs105225151*) is associated with the TRPV2 protein. (D) Box plot showing the expression levels between reference allele and the alternative allele. (E) Diagram illustrating a causal relationship between a genotype, TRPV2 expression level, and a behavioral trait.

We further explored the link between 464 proteins with *cis*-pQTL and 234 phenotypic traits in GeneNetwork.org^36–38^ that were previously collected for the HXB/BXH family. Among these 234 phenotypic traits, 68 traits showed significantly associated with genomic loci (LRS > 13.8; LOD > 3). Using the coloc program for colocalization analysis, we found two proteins with significant *cis*-pQTLs that colocalized with phenotypic QTLs. Notably, one of these signals shared the same leading QTL. This QTL (i.e., SNP: *rs107228843*; Chr4: 11,422,002 bp) was found to be associated with a phenotype of basal glucose uptake (GN trait ID: 10082) (**Figure S8A**), with a mapping LRS of 21.95. Another SNP (i.e., SNP: *D4Rat7*; Chr4: 11,782,356 bp) was found to be associated with the expression level of N-acyl phosphatidylethanolamine phospholipase D (NAPEPLD), with a mapping *p* value of 1.19 ×10^−6^ (**Figure S8B**). The reference allele has 2.84-fold (18.24 vs 16.82) higher expression than the alternative allele (**Figure S8A inset**). A causality analysis revealed that this locus potentially has a pleiotropic effect on both protein expression and phenotype (**Figure S8C**) using the Bayesian Network Webserver (BNW) program^51,52^.

### Pangenome analysis improves the identification of causal variants in QTLs

One commonly employed approach for identifying candidate genes within a QTL interval involves the identification of functional variants between two parental strains in a mapping population, such as the BXH/HXB panel. However, the approach is typically limited in resolution as it relies on mapping markers from the two parent strains. A better alternative is to leverage the pan-genome, which encompasses genome sequences for all HXB/BXH RI strains, to explore the variation across all mapping strains. To demonstrate candidate gene discovery using the pangenome, we select a small subset of proteins (6 proteins with *cis*-pQTL believed to influence phenotypic traits), for pangenome analysis. Among these, Fah (Fumarylacetoacetate Hydrolase) emerged as an intriguing example. Fah is mapped as a *cis*-pQTL with a mapping *p*-value of 6.53 × 10^−5^ (**Figure S9B**). The QTL is colocalized with a phenotypic QTL modulating cysteine level (**Figure S9A**). The expression level of Fah exhibited a high correlation with the levels of plasma cysteine (**Figure S9C**). To investigate functional variants within the Fah gene, we utilized PanGenome Graph Builder^53^ to obtain an unbiased graph representation^54^ of the pangenome of this gene and then applied ODGI ^55^ to understand its variation in the HXB/BXH family. Our analysis revealed a 142 bp insertion and deletion (INDEL) across the BXH/HXB strains (**Figure S9D, E**). The INDEL pattern is concordant with the expression of Fah across the strains. The rat pangenome enhances the discovery of candidate functional variants associated with the pQTLs and phenotypic variations.

### Genetic regulation of protein expression in rat point to human diseases

Rats, sharing extensive physiological and genetic similarities with humans, have served as a valuable model for translating protein expression variation insights from rodent studies to human conditions. We attempted to establish links between proteins with pQTL and human disorders (**Figure S10**). Using alcohol addiction as an example, we extracted 255 risk genes that are associated with “Ever addicted to alcohol” from the PhenomeXcan database ^56^. Among these, one protein (GNAO1) was found to be *trans*-regulated by a locus (*rs64148360*) with a mapping *p*-value of 7.97 × 10^−6^. GNAO1, a G protein subunit alpha o1, mediates the function of various neuronal receptors, including those implicated in alcohol addiction. Previous studies demonstrated an increase in GNAO protein expression levels in the rat cerebral cortex and cerebellum in response to chronic alcohol consumption ^57^. This analysis highlights the translational potential of rat models in unraveling molecular mechanisms relevant to human disorders.

## Discussion

In the present study, we aimed to elucidate the intricate mechanisms governing genetic modulation of protein expression in the rat brain and to understand its subsequent effects on phenotypic traits. To this end, we conducted a pilot proteome profiling of the brain across 29 rat HXB/BXH RI strains, as well as their two parental strains. This analysis enabled us to delineate the genetic modulation landscape of protein expression in the rat brain. We investigated the association between pQTLs and behavioral traits and used pangenome analysis to identify potential genetic variants for pQTLs and phenotypic QTLs. Moreover, we explored the potential link between rat pQTLs and human disorders. Collectively, these findings contribute to our understanding of the genetic regulation of protein expression in the rat brain and its potential link to phenotypic variation.

Recent research has consistently demonstrated that variations in both transcript-level expression and phenotypic traits tend to be influenced by noncoding variants^58^. Our present study revealed that the protein expression level tends to be regulated by non-synonymous variants. Our analysis revealed that 4.98% of *cis*-pQTLs reside in exonic regions, which is higher compared to the 2.85% of *cis*-eQTLs residing in exonic regions (**Figure S11**). This observation is consistent with our findings in the human pQTL analysis ^24^, underscoring the higher prevalence of pQTLs in exonic regions. These results suggest the involvement of potential post-transcriptional regulatory mechanisms in the control of protein expression. In addition, the number of pQTLs appears to be lower than that of eQTLs, indicating that a potential genetic buffering effect on the regulation of protein expression.

We have provided what we believe to be the most comprehensive proteomic data for the HXB/BXH rat RI strains to date. This dataset offers a valuable resource to explore the variation in sequence, expression, and pathways between SHR and BN-Lx strains. Employing a proteogenomics approach, we have successfully identified 97 variant peptides carrying non-synonymous variants. This number is comparable to the findings reported in previous studies ^35^. In addition, we observed that the pathway associated with mitochondrial reactive oxygen species (ROS) is highly enriched in the DEPs between SHR and BN-Lx strains at the protein expression level, which is consistent with the previous finding that genomic variants between rat substrains, including SHR and BN-Lx, are also enriched in the ROS pathway ^59^.

The pQTL analysis in this study was conducted using a relatively modest number of strains (*n* = 29). This number is small, especially when compared to human studies and other rodent models, such as collaborative mouse crosses. Due to this small sample size, our study may face limitations in statistical power, potentially impacting the number of detected pQTLs. However, we envision significant improvements in our ability to capture a broader range of pQTLs with the inclusion of additional RI strains, such as the FXLE/LEXF panel. By expanding the diversity of strains analyzed, we aim to achieve a more comprehensive understanding of the genetic modulation of protein expression in the rat brain, overcoming the current limitations associated with the sample size. The expansion holds promise for unveiling a richer landscape of regulatory genetic elements influencing protein expression.

This is one of the first studies that applies a rat pangenome in genetics. It is also one of the first studies to take pQTL and successfully use the pangenome to zoom in on underlying variants displaying genetic variants within the Fah gene, with an evident large insertion highlighted in SHR and some HXB strains that was never seen before. The emerging field of pangenomics allows the study of full genomes where the genome of all strains can be directly compared to each other. It provides the lossless comparison of all (complex) variants between individuals directly without the use of a single reference genome and inherited reference bias^60,61^. This pangenomic approach will be especially powerful when individual genomes are all assembled from long-read sequence data, some of which are already available for the upcoming rat pangenome. This will enable a complete catalog of all genomic variants, including SVs and repeats, that differ between individuals and will be beneficial to future protein studies in rat.

In conclusion, our study offers a large-scale protein expression profile in the rat brain. We defined the genetic regulation of protein expression in the brain, emphasizing a wide range of exonic variants. We have successfully established causal links between identified protein pQTLs and phenotypic traits. Investigation of the possible connection between rat pQTLs and human disorders revealed insights that can inform translational studies, bridging the gap between animal models and human health. By elucidating the mechanisms of protein expression regulation, our work opens new pathways for discovering novel candidate genes associated with diverse phenotypes, contributing significantly to the field of genetics and neurobiology.

## Online methods

### Animals

For this study, we utilized both male and female rats aged two months, from a total of 29 HXB/BXH RI strains and two parental strains. All experimental procedures involving animals were carried out in strict adherence to the guidelines approved by the Institutional Ethics Committees of the Institute of Physiology, Czech Academy of Sciences, located in Prague, Czech Republic.

### Behavioral phenotyping

Male rats of the SHR/Ola and BN-Lx/Cub progenitor strains and 30 RI strains (being inbred for more than 90 generations) were obtained from the Department of Genetics of Model Diseases at the Institute of Physiology, Academy of Sciences of the Czech Republic. Number of rats in each strain ranged from 6 to 14. Totally, we used 277 animals. The animals were housed in an air-conditioned animal room with a stable temperature (22oC), humidity (40%) and 12/12 light/dark cycle (light on at 7:00 am). Animals were accommodated in the animal room for two weeks and behavioral testing was done at the age of 11-14 weeks. For testing in the active place avoidance task, requiring aversive reinforcement, conscious animals were gently implanted with a subcutaneous needle, which pierced the rat’s skin between its shoulders and its sharp end was cut and bent to form a small loop, preventing slip-out and providing purchase for an alligator clip delivering shocks. This procedure loads minimal stress onto animals and is analogous to subcutaneous injection in humans. Water and food were freely available to the rats throughout the study. All experimentation complied with current legislation on protection of laboratory animals (Animal Protection Code of the Czech Republic, European Union directives 2010/63/EC; 86/609/EEC and NIH guidelines).

The apparatus has been described previously in detail ^62–64^. Briefly, it consisted of a smooth metal disk (82 cm in diameter; elevated 1 m above the room floor), which rotated clockwise at 1 revolution per minute (rpm). A 50-cm-high transparent wall made of clear Plexiglas surrounded the arena. The animals had to avoid an unmarked, 60-deg to-be-avoided sector identified solely by its relationships to distal room cues. The position of the shock sector remained stable in the throughout each phase. The only change of the sector position occurred before the reversal phase.

The rats again wore a rubber jacket, which carried an infrared LED between animal’s shoulders. A computer-based tracking system (iTrack; Tracker, Biosignal Group, USA) located in an adjacent room recorded the rat’s position at a frequency of 25 Hz. Position series were stored for off-line analysis (TrackAnalysis; Biosignal Group, USA). Whenever the rat entered the to-be-avoided sector for more than 300 ms, the tracking system delivered a mild electric shock (50 Hz, 0.5 s, 0.4-0.6 mA) and counted an entrance. If the rat did not leave the sector, additional shocks were given every 900 ms, but no more entrances were counted until the rat left the sector for more than 300 ms. Shocks were delivered through the implanted needle and the grounded arena floor (the highest voltage drop, and perception of shocks were between rats’ paws and floor). This shocking procedure was previously shown to be effective and safe for rats, leading to rapid avoidance behavior ^62,65–67^. The current was individualized for each rat to elicit a rapid escape response but to prevent freezing, however, in most cases, animals responded appropriately to 0.4 mA. There were no systematic differences between shock intensities between strains. After each rat, the floor was cleaned with ethanol, ensuring the rats could not use inter-trial scent marks. Rats were tested in semi-mixed design to avoid batch effects.

### Protein extraction, quantification, and digestion

Whole rat brain tissues were weighed and homogenized using a lysis buffer composed of 50 mM HEPES (pH 8.5), 8 M urea, and 0.5% sodium deoxycholate, with a ratio of 100 µl of buffer per 10 mg of tissue. To inhibit phosphatase activity, a 1×PhosSTOP phosphatase inhibitor cocktail (Sigma-Aldrich) was added to the lysis buffer. The total protein concentration of each sample was determined using the BCA Protein Assay Kit (Thermo Fisher Scientific) and further confirmed through Coomassie staining of short SDS gels ^68,69^. Quantified protein samples (∼0.3 mg) in the lysis buffer containing 8 M urea were first subjected to proteolysis using Lys-C (Wako) at a ratio of 1:100 (enzyme to protein) at room temperature for a duration of 2 hours. Subsequently, the samples were diluted four-fold to reduce the urea concentration to 2 M, followed by digestion with trypsin (Promega) at a ratio of 1:50 (enzyme to protein) at room temperature overnight. To terminate the digestion process, 1% trifluoroacetic acid was added, and the mixture was then centrifuged.

### Tandem mass tag labeling

Following the completion of the digestion and termination steps, the supernatant was subjected to desalting using a Sep-Pak C18 cartridge (Waters), and subsequently dried using a speedvac. Each individual sample was then reconstituted in 50 mM HEPES (pH 8.5) and labeled with either 11-plex or 16-plex TMT reagents. These labeled samples were mixed in equal proportions and subjected to another round of desalting in preparation for subsequent fractionation. A protein amount of 0.1 mg was used for each sample. The overall experimental design included a total of 2 batches of 11-plex TMT experiments and 3 batches of 16-plex TMT experiments.

### Extensive two-dimensional LC/LC-MS/MS

The pooled samples, labeled with TMT, were subjected to fractionation using offline basic pH reversed-phase chromatography (HPLC), followed by acidic pH reverse phase LC-MS/MS analysis. The offline basic HPLC was conducted according to previously published methods ^70,71^. In the 16-plex TMT batch, we generated 40 concatenated fractions, while in the 11-plex TMT batch, 36 concatenated fractions were generated. The offline LC run was carried out using an XBridge C_18_ column (3.5 μm particle size, 4.6 mm x 25 cm, Waters) with a 3-hour gradient. The mobile phase consisted of buffer A (10 mM ammonium formate, pH 8.0) and buffer B (95% acetonitrile, 10 mM ammonium formate, pH 8.0)^68^.

For the subsequent acidic pH LC-MS/MS analysis, each fraction was sequentially run on a column (75 µm x 15-30 cm, 1.9 µm C18 resin from Dr. Maisch GmbH) heated to 65° C to reduce backpressure. The column was interfaced with an Orbitrap Fusion and Q Exactive HF MS system (Thermo Fisher). Peptides were eluted using a 1.5-2-hour gradient with buffer A (0.2% formic acid, 5% DMSO) and buffer B (buffer A supplemented with 65% acetonitrile). The MS settings included MS1 scans with a resolution of 60,000, an AGC target of 1 x 10^6^, and a maximal ion time of 100 ms. Additionally, 20 data-dependent MS2 scans were performed in the m/z range of 410-1600, with a resolution of 60,000, an AGC target of 1 x 10^5^, a maximal ion time of ∼10^5^ ms, and higher-energy collisional dissociation (HCD) with 38% normalized collision energy. The isolation window was set to 1.0 m/z with a 0.2 m/z offset, and dynamic exclusion was applied for approximately 15 seconds.

### Identification of proteins by database search with JUMP software

To improve the sensitivity and specificity, we performed peptide identification with the JUMP search engine ^72^. JUMP searched MS/MS raw data against a composite target/decoy database ^73^ to evaluate FDR. The target rat protein sequences (29,940 entries) were downloaded from the UniProt database. The decoy database, generated by reversing the target sequences, was concatenated to the target database. FDR was estimated by calculating the ratio of the number of decoy matches to the number of target matches. Major parameters included precursor and product ion mass tolerance (± 15 ppm), full trypticity, static mass shift for the TMT tags (+229.16293/+304.20715) and carbamidomethyl modification of 57.02146 on cysteine, dynamic mass shift for Met oxidation (+15.99491), maximal missed cleavage (*n* = 2), and maximal modification sites (*n* = 3). Putative PSMs were filtered by mass accuracy and then grouped by precursor ion charge state and filtered by JUMP-based matching scores (Jscore and ΔJn) to reduce protein FDR below 1%. If one peptide could be generated from multiple homologous proteins, based on the rule of parsimony, the peptide was assigned to the canonical protein form in the manually curated Swiss-Prot database.

### Protein quantification by JUMP software suite

Protein quantification was carried out using the following steps ^74^. We first extracted TMT reporter ion intensities of each PSM and corrected the raw intensities based on isotopic distribution of each labeling reagent. We discarded PSMs with low intensities (i.e., the minimum intensity of 1,000 and median intensity of 5,000). After normalizing abundance with the trimmed median intensity of all PSMs, we calculated the mean-centered intensities across samples (e.g., relative intensities between each sample and the mean) and summarized protein relative intensities by averaging related PSMs. Finally, we derived protein absolute intensities by multiplying the relative intensities by the grand mean of the three most highly abundant PSMs. Log_2_-transformed data were used for the subsequent normalized analysis.

### Proteogenomic analysis

Sequence variants between SHR and BN-Lx strains were provided by the rat reference genome project ^42^. All variants were re-annotated with the ANNOVAR program^75^. A total of 8,723 missense variants were extracted for the proteogenomics analysis. We used JUMPg, a proteogenomics pipeline, to detect variant peptides.

### RNA-seq data analysis

Transcriptomics data were obtained from a previous study ^40^, which included a total of 94 samples from 30 HXB/BXH recombinant inbred (RI) strains, as well as their parental strains, SHR and BN-Lx. The dataset consisted of expression levels for 18,385 genes and included 2-3 replicates per strain. To facilitate subsequent analysis, the mean expression values for each strain were calculated and recorded.

### Genotype data

Genotypes of the HXB/BXH were downloaded from the Genenetwork database ^76^. The genotypes were annotated with the Rnor6.0 genomic assembly ^42^. A total of 10,465 genotypes from 30 rat strains were used for the QTL mapping.

### Principal component analysis

Principal component analysis (PCA) was used to visualize the differences among samples. All gene and metabolite abundance were used as features of PCA. The pairwise Euclidean distance between features was calculated. PCA was performed using the R package prcomp (version 3.4.0)^77^.

### Differential expression analysis

Differentially expressed proteins between the two strains were identified using the limma R package (version 3.46.0)^39^. The Benjamini-Hochberg method for false discovery rate correction was used, and proteins with an adjusted *p* value < 0.01 and fold change > 2 were defined as differentially expressed between the SHR and BN-Lx parental strains.

### Pathway enrichment

To assess the functional relevance of the differentially expressed proteins, the R package clusterProfiler (version 3.18.1) ^78^ was used for Kyoto Encyclopedia of Genes and Genomes (KEGG) enrichment analysis. KEGG terms with a Benjamini-Hochberg *p* value < 0.05 were defined as significantly enriched.

### Analysis of the heritability of protein expression

Heritability of protein expression was determined by assessing the proportion of the total variance that can be attributed to additive genetic effects. The total variance was computed as the sum of squared differences from the mean expression for all 62 samples, whereas the additive variance was calculated by summing the square differences from the mean expression for all 31 rat strains after averaging data from two biological replicates (i.e., male and female for each strain).

### Linkage analysis

For each protein, we estimated the proportion of variance in protein expression level explained by all SNPs using a linear mixed model (LMM) in GEMMA ^79^ that corrects for kinship. GEMMA is the primary mapper in GeneNetwork.org ^36–38^. The top variant was selected as the QTL for the protein/gene eQTLs/pQTLs were defined as *cis* (local) if the QTL was within 1 Mb on either side of the TSS, whereas eQTLs/pQTLs were defined as *trans* (distal) if the peak association was at least 5 Mb outside of the exon boundaries^79,80^.

We permuted the sample labels ten times and applied the same pQTL mapping procedure to obtain an empirical null distribution of gene-level *p* values^81–83^. After each permutation, we kept the most significant *p* value for each protein. With the empirical null distribution, we computed the false discovery rate (FDR) associated with each *p* value threshold and selected the p value threshold that provided a 5% FDR control.

### Bayesian network analysis

We examined the causal connections among genotypes, protein expression levels, and phenotypes using the Bayesian network modeling tool (BNW, compbio.uthsc.edu/BNW) ^51,52^. In this study, the BNW on GeneNetwork was employed with genotypes linked to a specific phenotype, proteins exhibiting pQTLs within the phenotypic QTL region, and phenotypic data as input variables. The analysis tests the causal dependencies between genetic variations, protein abundance, and phenotypic traits using Bayesian network.

### Functional annotation of QTLs

ANNOVAR^75^ was used to functionally annotate the leading SNP of a QTL. RefSeq from the UCSC genome browser database was used to annotate SNPs. The functional consequence (synonymous, non-synonymous) of coding SNP was also determined.

### Co-localization analysis

We utilized the coloc R package^84^ to assess the colocalization signals between *cis*-eQTLs and *cis*-pQTLs. A window of 500 kb was applied on both flanks of the pQTL locus. The coloc program was employed to derive posterior probabilities for five distinct yet mutually exclusive hypotheses. These hypotheses include: (i) no connection between any variant within the region and either *cis*-pQTL or *cis*-eQTL (H0); (ii) exclusive linkage with *cis*-pQTL excluding *cis*-eQTL (H1); (iii) exclusive linkage with *cis*-eQTL excluding *cis*-pQTL (H2); (iv) presence of two distinct QTLs (H3); and (v) presence of a shared QTL influencing both gene and protein expression (H4). A PP4 value exceeding 0.8 was considered a significant indication of colocalization.

### Zooming in on variants using the rat pangenome

Gene pangenome graphs were built using the Pangenome Graph Builder (PGGB) ^53^ that include wfmash alignments ^85^, and SEQWISH ^54^ to produce unbiased graph models that losslessly represent both sequences and their variation. We then used ODGI ^55^, VG ^86^ and Bandage ^87^ to analyze the graph and produce pangenome visualizations that helped us zoom in gene candidates and structural variants

## Acknowledgments

This work was supported by NIH/NIDA P30 Pilot Grant (1P30DA044223). L.L. and X.W. were also partially supported by NIH/NIA R01 (RF1AG072703) and NIH/NIDA R01 (R01DA056523). M.P. supported by the National Institute for Research of Metabolic and Cardiovascular Diseases (Program EXCELES, ID Project No. LX22NPO5104) funded by the European Union – Next Generation E.U., H.B., K.V., and A.S. were supported in part by the Czech Science Foundation (GACR) grants (21-16667K) and by the project National Institute for Neurology Research (Programme EXCELES, ID Project No. LX22NPO5107) Funded by the European Union–Next Generation EU.

## Author Contributions

XW, JP, and RW planned and supervised all experiments and data analysis. LL, FV, AG, DK, AZ and AG provided bioinformatics support and carried out data analysis. ZW and AM performed TMT-based proteomics experiment. LS and HC provided transcriptomic and genomic variant data, respectively. HB, KV, AS and MP conducted the behavioral testing and analyzed the results. LL, RW, and XW wrote the manuscript.

## Declaration of Interests

The authors declare no competing interests.

## Data availability

The mass spectrometry proteomics data have been deposited to the Proteome Xchange Consortium via the PRIDE ^88^ partner repository with the data set identifier PXD048286.

## Code availability

Data analyses were performed in LINUX shell, Perl (v5.18.4), and R (v4.0.4; https://www.r-project.org/). Annotations were performed using ANNOVAR (https://annovar.openbioinformatics.org/en/latest/user-guide/startup/). Proteomic data were processed with JUMP software (https://github.com/JUMPSuite/JUMP), and QTL mapping was performed using GEMMA (https://github.com/genetics-statistics/GEMMA) and coloc (https://github.com/cran/coloc). The STRING database was utilized for network analysis (https://string-db.org/). All pangenome tools are available through https://github.com/pangenome/

## Notes

### Competing Interest Statement

The authors have declared no competing interest.

### Summary of Updates

The authors list has been modified.

## References

1. Jacob HJ, Brown DM, Bunker RK, et al. A genetic linkage map of the laboratory rat, Rattus norvegicus. Nat Genet. 1995;9(1):63–69.

2. Tao YT, Ding XB, Jin J, et al. Predicted rat interactome database and gene set linkage analysis. Database (Oxford). 2020;2020.

3. Szpirer C. Rat models of human diseases and related phenotypes: a systematic inventory of the causative genes. J Biomed Sci. 2020;27(1):84.

4. Civelek M, Lusis AJ. Systems genetics approaches to understand complex traits. Nat Rev Genet. 2014;15(1):34–48.

5. Chen J, Zhao X, Cui L, et al. Genetic regulatory subnetworks and key regulating genes in rat hippocampus perturbed by prenatal malnutrition: implications for major brain disorders. Aging (Albany NY). 2020;12(9):8434–8458.

6. Yang J, Hu C, Hu H, et al. QTLNetwork: mapping and visualizing genetic architecture of complex traits in experimental populations. Bioinformatics. 2008;24(5):721–723.

7. Ashbrook DG, Arends D, Prins P, et al. A platform for experimental precision medicine: The extended BXD mouse family. Cell Syst. 2021;12(3):235–247 e239.

8. Trotter C, Kim H, Farage G, et al. Speeding up eQTL scans in the BXD population using GPUs. G3 (Bethesda). 2021;11(12).

9. Gusev A, Mancuso N, Won H, et al. Transcriptome-wide association study of schizophrenia and chromatin activity yields mechanistic disease insights. Nat Genet. 2018;50(4):538–548.

10. Wang D, Liu S, Warrell J, et al. Comprehensive functional genomic resource and integrative model for the human brain. Science. 2018;362(6420).

11. Wu L, Candille SI, Choi Y, et al. Variation and genetic control of protein abundance in humans. Nature. 2013;499(7456):79–82.

12. Sun BB, Maranville JC, Peters JE, et al. Genomic atlas of the human plasma proteome. Nature. 2018;558(7708):73–79.

13. Yang C, Farias FHG, Ibanez L, et al. Genomic atlas of the proteome from brain, CSF and plasma prioritizes proteins implicated in neurological disorders. Nat Neurosci. 2021;24(9):1302–1312.

14. Wingo TS, Liu Y, Gerasimov ES, et al. Brain proteome-wide association study implicates novel proteins in depression pathogenesis. Nat Neurosci. 2021;24(6):810–817.

15. Suhre K, McCarthy MI, Schwenk JM. Genetics meets proteomics: perspectives for large population-based studies. Nat Rev Genet. 2021;22(1):19–37.

16. Li L, Niu MM, Erickson A, et al. SMAP is a pipeline for sample matching in proteogenomics. Nat Commun. 2022;13(1).

17. Wu Y, Williams EG, Dubuis S, et al. Multilayered genetic and omics dissection of mitochondrial activity in a mouse reference population. Cell. 2014;158(6):1415–1430.

18. Williams EG, Wu Y, Jha P, et al. Systems proteomics of liver mitochondria function. Science. 2016;352(6291):aad0189.

19. Williams EG, Pfister N, Roy S, et al. Multiomic profiling of the liver across diets and age in a diverse mouse population. Cell Syst. 2022;13(1):43–57 e46.

20. Chick JM, Munger SC, Simecek P, et al. Defining the consequences of genetic variation on a proteome-wide scale. Nature. 2016;534(7608):500–505.

21. Keele GR, Zhang T, Pham DT, et al. Regulation of protein abundance in genetically diverse mouse populations. Cell Genom. 2021;1(1).

22. Hubner N, Wallace CA, Zimdahl H, et al. Integrated transcriptional profiling and linkage analysis for identification of genes underlying disease. Nat Genet. 2005;37(3):243–253.

23. Printz MP, Jirout M, Jaworski R, Alemayehu A, Kren V. Genetic Models in Applied Physiology. HXB/BXH rat recombinant inbred strain platform: a newly enhanced tool for cardiovascular, behavioral, and developmental genetics and genomics. J Appl Physiol (1985). 2003;94(6):2510–2522.

24. Luo J, Niu M, Li L, et al. Genetic regulation of human brain proteome reveals proteins implicated in psychiatric disorders. Research Square. 2022.

25. Adriaens ME, Lodder EM, Moreno-Moral A, et al. Systems Genetics Approaches in Rat Identify Novel Genes and Gene Networks Associated With Cardiac Conduction. J Am Heart Assoc. 2018;7(21):e009243.

26. Aitman TJ, Critser JK, Cuppen E, et al. Progress and prospects in rat genetics: a community view. Nat Genet. 2008;40(5):516–522.

27. Atanur SS, Diaz AG, Maratou K, et al. Genome sequencing reveals loci under artificial selection that underlie disease phenotypes in the laboratory rat. Cell. 2013;154(3):691–703.

28. Tabakoff B, Saba L, Printz M, et al. Genetical genomic determinants of alcohol consumption in rats and humans. BMC Biol. 2009;7:70.

29. Lusk R, Hoffman PL, Mahaffey S, et al. Beyond Genes: Inclusion of Alternative Splicing and Alternative Polyadenylation to Assess the Genetic Architecture of Predisposition to Voluntary Alcohol Consumption in Brain of the HXB/BXH Recombinant Inbred Rat Panel. Front Genet. 2022;13:821026.

30. Saba LM, Hoffman PL, Homanics GE, et al. A long non-coding RNA (Lrap) modulates brain gene expression and levels of alcohol consumption in rats. Genes Brain Behav. 2021;20(2):e12698.

31. Simonis M, Atanur SS, Linsen S, et al. Genetic basis of transcriptome differences between the founder strains of the rat HXB/BXH recombinant inbred panel. Genome Biol. 2012;13(4):r31.

32. Senko AN, Overall RW, Silhavy J, et al. Systems genetics in the rat HXB/BXH family identifies Tti2 as a pleiotropic quantitative trait gene for adult hippocampal neurogenesis and serum glucose. Plos Genet. 2022;18(4).

33. Bettenworth D, Bokemeyer A, Baker M, et al. Assessment of Crohn’s disease-associated small bowel strictures and fibrosis on cross-sectional imaging: a systematic review. Gut. 2019;68(6):1115–1126.

34. Gordon IO, Bettenworth D, Bokemeyer A, et al. Histopathology Scoring Systems of Stenosis Associated With Small Bowel Crohn’s Disease: A Systematic Review. Gastroenterology. 2020;158(1):137–150 e131.

35. Low TY, van Heesch S, van den Toorn H, et al. Quantitative and qualitative proteome characteristics extracted from in-depth integrated genomics and proteomics analysis. Cell Rep. 2013;5(5):1469–1478.

36. Mulligan MK, Mozhui K, Prins P, Williams RW. GeneNetwork: A Toolbox for Systems Genetics. Methods Mol Biol. 2017;1488:75–120.

37. Anderson KR, Harris JA, Ng L, et al. Highlights from the Era of Open Source Web-Based Tools. J Neurosci. 2021;41(5):927–936.

38. Sloan Z, Arends D, W. Broman K, et al. GeneNetwork: framework for web-based genetics. The Journal of Open Source Software. 2016;1(2).

39. Ritchie ME, Phipson B, Wu D, et al. limma powers differential expression analyses for RNA-sequencing and microarray studies. Nucleic Acids Res. 2015;43(7):e47.

40. Saba LM, Flink SC, Vanderlinden LA, et al. The sequenced rat brain transcriptome--its use in identifying networks predisposing alcohol consumption. FEBS J. 2015;282(18):3556–3578.

41. Li Y, Wang X, Cho JH, et al. JUMPg: An Integrative Proteogenomics Pipeline Identifying Unannotated Proteins in Human Brain and Cancer Cells. J Proteome Res. 2016;15(7):2309–2320.

42. de Jong TV, Pan Y, Rastas P, et al. A revamped rat reference genome improves the discovery of genetic diversity in laboratory rats. bioRxiv. 2023.

43. Morrissey C, Grieve IC, Heinig M, et al. Integrated genomic approaches to identification of candidate genes underlying metabolic and cardiovascular phenotypes in the spontaneously hypertensive rat. Physiol Genomics. 2011;43(21):1207–1218.

44. Cheng Y, Sun D, Zhu B, et al. Integrative Metabolic and Proteomic Profiling of the Brainstem in Spontaneously Hypertensive Rats. J Proteome Res. 2020;19(10):4114–4124.

45. Chen Y, Wu L, Liu J, Ma L, Zhang W. Adenine nucleotide translocase: Current knowledge in post-translational modifications, regulations and pathological implications for human diseases. FASEB J. 2023;37(6):e22953.

46. Alnouti Y, Klaassen CD. Tissue distribution, ontogeny, and regulation of aldehyde dehydrogenase (Aldh) enzymes mRNA by prototypical microsomal enzyme inducers in mice. Toxicol Sci. 2008;101(1):51–64.

47. Vasiliou V, Pappa A, Petersen DR. Role of aldehyde dehydrogenases in endogenous and xenobiotic metabolism. Chem Biol Interact. 2000;129(1-2):1–19.

48. Singh S, Brocker C, Koppaka V, et al. Aldehyde dehydrogenases in cellular responses to oxidative/electrophilic stress. Free Radic Biol Med. 2013;56:89–101.

49. Marchitti SA, Brocker C, Stagos D, Vasiliou V. Non-P450 aldehyde oxidizing enzymes: the aldehyde dehydrogenase superfamily. Expert Opin Drug Metab Toxicol. 2008;4(6):697–720.

50. Rotondo F, Ho-Palma AC, Remesar X, Fernandez-Lopez JA, Romero MDM, Alemany M. Effect of sex on glucose handling by adipocytes isolated from rat subcutaneous, mesenteric and perigonadal adipose tissue. PeerJ. 2018;6:e5440.

51. Ziebarth JD, Bhattacharya A, Cui Y. Bayesian Network Webserver: a comprehensive tool for biological network modeling. Bioinformatics. 2013;29(21):2801–2803.

52. Ziebarth JD, Cui Y. Precise Network Modeling of Systems Genetics Data Using the Bayesian Network Webserver. Methods Mol Biol. 2017;1488:319–335.

53. Erik Garrison AG, Simon Heumos, Flavia Villani, Zhigui Bao, Lorenzo Tattini, Jörg Hagmann, Sebastian Vorbrugg, Santiago Marco-Sola, Christian Kubica, David G. Ashbrook, Kaisa Thorell, Rachel L. Rusholme-Pilcher, Gianni Liti, Emilio Rudbe, Sven Nahnsen, Zuyu Yang, Mwaniki N. Moses, Franklin L. Nobrega, Yi Wu, Hao Chen, Joep de Ligt, Peter H. Sudmant, Nicole Soranzo, Vincenza Colonna, Robert W. Williams, Pjotr Prins. Building pangenome graphs. bioRxiv.2023.2004.2005.535718.

54. Garrison E, Guarracino A. Unbiased pangenome graphs. Bioinformatics. 2023;39(1).

55. Guarracino A, Heumos S, Nahnsen S, Prins P, Garrison E. ODGI: understanding pangenome graphs. Bioinformatics. 2022;38(13):3319–3326.

56. Pividori M, Rajagopal PS, Barbeira A, et al. PhenomeXcan: Mapping the genome to the phenome through the transcriptome. Sci Adv. 2020;6(37).

57. Guillen A, Garcia-Villafranca J, Mata F, Homburger V, Castro J. Effect of chronic alcohol consumption on Go protein in rat brain. Neurosci Lett. 2003;353(3):177–180.

58. Munro D, Wang T, Chitre AS, et al. The regulatory landscape of multiple brain regions in outbred heterogeneous stock rats. Nucleic Acids Res. 2022;50(19):10882–10895.

59. Hermsen R, de Ligt J, Spee W, et al. Genomic landscape of rat strain and substrain variation. BMC Genomics. 2015;16(1):357.

60. Liao WW, Asri M, Ebler J, et al. A draft human pangenome reference. Nature. 2023;617(7960):312–324.

61. Guarracino A, Buonaiuto S, de Lima LG, et al. Recombination between heterologous human acrocentric chromosomes. Nature. 2023;617(7960):335–343.

62. Wesierska M, Dockery C, Fenton AA. Beyond memory, navigation, and inhibition: behavioral evidence for hippocampus-dependent cognitive coordination in the rat. J Neurosci. 2005;25(9):2413–2419.

63. Kubik S, Fenton AA. Behavioral evidence that segregation and representation are dissociable hippocampal functions. J Neurosci. 2005;25(40):9205–9212.

64. Stuchlik A, Petrasek T, Prokopova I, et al. Place avoidance tasks as tools in the behavioral neuroscience of learning and memory. Physiol Res. 2013;62(Suppl 1):S1–S19.

65. Š. Kubík AS, A.A. FENTON1. Evidence for hippocampal role in place avoidance other than merely memory storage. Physiol Res. 2006;55.

66. Stuchlik A, Petrasek T, Vales K. Dopamine D2 receptors and alpha1-adrenoceptors synergistically modulate locomotion and behavior of rats in a place avoidance task. Behav Brain Res. 2008;189(1):139–144.

67. Stuchlik A, Vales K. Baclofen dose-dependently disrupts learning in a place avoidance task requiring cognitive coordination. Physiol Behav. 2009;97(3-4):507–511.

68. Bai B, Tan H, Pagala VR, et al. Deep Profiling of Proteome and Phosphoproteome by Isobaric Labeling, Extensive Liquid Chromatography, and Mass Spectrometry. Methods Enzymol. 2017;585:377–395.

69. Pagala VR, High AA, Wang X, et al. Quantitative protein analysis by mass spectrometry. Methods Mol Biol. 2015;1278:281–305.

70. Xu P, Duong DM, Peng J. Systematical optimization of reverse-phase chromatography for shotgun proteomics. J Proteome Res. 2009;8(8):3944–3950.

71. Wang H, Yang Y, Li Y, et al. Systematic optimization of long gradient chromatography mass spectrometry for deep analysis of brain proteome. J Proteome Res. 2015;14(2):829–838.

72. Wang X, Li Y, Wu Z, Wang H, Tan H, Peng J. JUMP: a tag-based database search tool for peptide identification with high sensitivity and accuracy. Mol Cell Proteomics. 2014;13(12):3663–3673.

73. Peng J, Elias JE, Thoreen CC, Licklider LJ, Gygi SP. Evaluation of multidimensional chromatography coupled with tandem mass spectrometry (LC/LC-MS/MS) for large-scale protein analysis: the yeast proteome. J Proteome Res. 2003;2(1):43–50.

74. Niu M, Cho JH, Kodali K, et al. Extensive Peptide Fractionation and y(1) Ion-Based Interference Detection Method for Enabling Accurate Quantification by Isobaric Labeling and Mass Spectrometry. Anal Chem. 2017;89(5):2956–2963.

75. Wang K, Li M, Hakonarson H. ANNOVAR: functional annotation of genetic variants from high-throughput sequencing data. Nucleic Acids Res. 2010;38(16):e164.

76. Jirout M, Krenova D, Kren V, et al. A new framework marker-based linkage map and SDPs for the rat HXB/BXH strain set. Mamm Genome. 2003;14(8):537–546.

77. Sadrara M, Khorrami MK. Principal component analysis-multivariate adaptive regression splines (PCA-MARS) and back propagation-artificial neural network (BP-ANN) methods for predicting the efficiency of oxidative desulfurization systems using ATR-FTIR spectroscopy. Spectrochim Acta A Mol Biomol Spectrosc. 2023;300:122944.

78. Yu GC, Wang LG, Han YY, He QY. clusterProfiler: an R Package for Comparing Biological Themes Among Gene Clusters. Omics. 2012;16(5):284–287.

79. Zhou X, Stephens M. Genome-wide efficient mixed-model analysis for association studies. Nat Genet. 2012;44(7):821–824.

80. Shang L, Smith JA, Zhao W, et al. Genetic Architecture of Gene Expression in European and African Americans: An eQTL Mapping Study in GENOA. Am J Hum Genet. 2020;106(4):496–512.

81. Jansen R, Hottenga JJ, Nivard MG, et al. Conditional eQTL analysis reveals allelic heterogeneity of gene expression. Hum Mol Genet. 2017;26(8):1444–1451.

82. Pickrell JK, Marioni JC, Pai AA, et al. Understanding mechanisms underlying human gene expression variation with RNA sequencing. Nature. 2010;464(7289):768–772.

83. Barreiro LB, Tailleux L, Pai AA, Gicquel B, Marioni JC, Gilad Y. Deciphering the genetic architecture of variation in the immune response to Mycobacterium tuberculosis infection. Proc Natl Acad Sci U S A. 2012;109(4):1204–1209.

84. Wang G, Sarkar A, Carbonetto P, Stephens M. A simple new approach to variable selection in regression, with application to genetic fine mapping. J R Stat Soc Series B Stat Methodol. 2020;82(5):1273–1300.

85. Guarracino AM, Njagi, Marco-Sola, Santiago, Garrison, Erik. wfmash: whole-chromosome pairwise alignment using the hierarchical wavefront algorithm (v0.10.4). Zenodo. 2023.

86. Hickey G, Heller D, Monlong J, et al. Genotyping structural variants in pangenome graphs using the vg toolkit. Genome Biol. 2020;21(1):35.

87. Wick RR, Schultz MB, Zobel J, Holt KE. Bandage: interactive visualization of de novo genome assemblies. Bioinformatics. 2015;31(20):3350–3352.

88. Deutsch EW, Bandeira N, Perez-Riverol Y, et al. The ProteomeXchange consortium at 10 years: 2023 update. Nucleic Acids Res. 2023;51(D1):D1539–D1548.

