## Supplemental Figures for "Genetic Modulation of Protein Expression in Rat Brain"

†Correspondence authors

Running title: Genetic architecture of rat brain proteome

**Supplementary Figures**

**Figure S1.** Workflow for the analysis of quantitative proteomics data.

**Figure S2.** Protein coverage and correlation analysis between two parental strains.

**Figure S3.** Comparison between proteomic and transcriptomic data.

**Figure** **S4**. Expression levels of variant peptides in SHR and BN-Lx strains.

**Figure S5.** Analysis of highly variable proteins across all 29 stains.

**Figure S6.** Determination of the *p*-value cutoff.

**Figure S7.** Sex-specific genetic regulation of the brain proteome.

**Figure S8.** Co-localization of *cis*-QTLs and phenotypic traits.

**Figure S9.** Pangenome analysis revealing genetic variation among mapping strains.

**Figure S10.** Manhattan plot showing a *trans*-pQTL is associated with GNAO1 protein expression.

**Figure S11.** Proportions of QTLs in different genomic regions.

**Supplementary Tables**

**Table S1**. TMT-LC/LC-MS/MS-based proteomic profile of whole brain tissues from HXB/BXH rat strains.

**Table S2A.** Variant peptides detected by JUMPg proteogenomics pipeline.

**Table S2B.** Differentially expressed proteins (DEPs) identified between the two parental strains of the HXB/BXH RI rat panel.

**Table S2C.** Enrichment analysis of proteins with higher expression in SHR compared to BN-Lx.

**Table S2D.** Enrichment analysis of proteins with higher expression in BN-Lx compared to SHR.

**Table S2E.** Enrichment analysis of proteins with variation coefficients.

**Table S3A**. pQTLs identified from sex-averaged expression levels of 29 RI strains.

**Table S3B**. Female-specific pQTLs identified in 29 RI strains.

**Table S3C.** Male-specific pQTLs identified in 29 RI strains.

**Table S4**. *cis*-eQTL identified in the rat brain tissue from 30 RI stains.

**Table S5**. Co-localization analysis for *cis*-pQTLs and *cis*-eQTLs.

**Table S6**. Co-localization analysis for *cis*-pQTLs and *trans*-pQTLs.

**Table S7**. Phenotypic traits used for the linkage analysis.

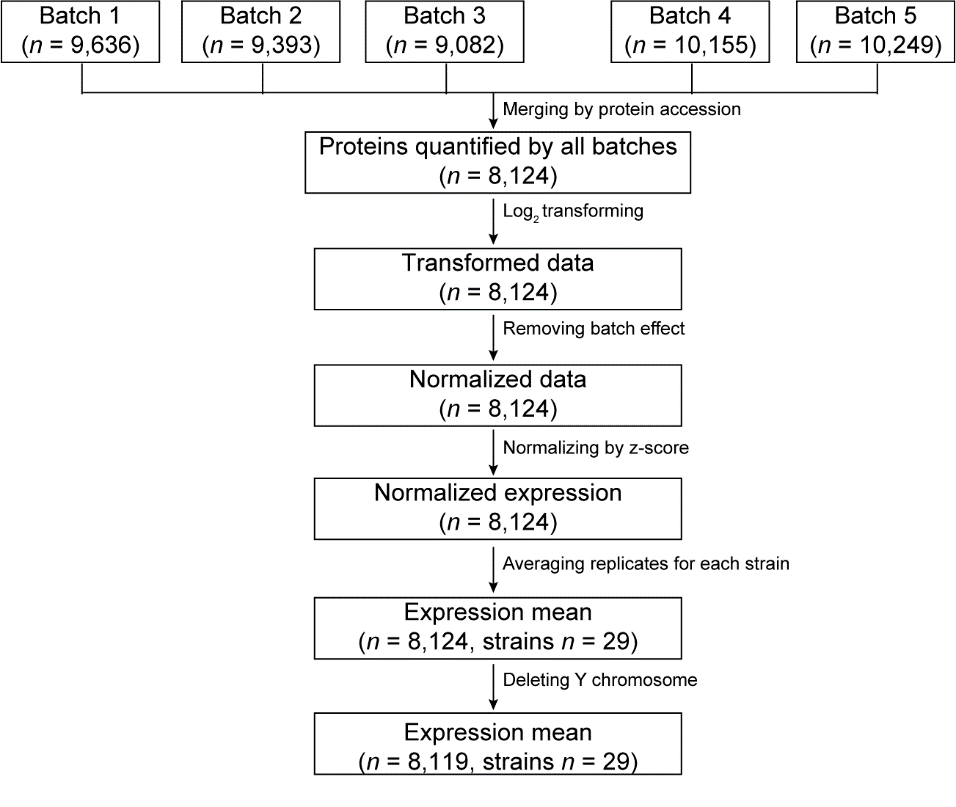

**Figure S1. Workflow for the analysis of quantitative proteomics data.** A total of five batches of TMT experiments were conducted, generating quantitative proteomics data for 62 samples. Proteins from each batch were merged based on their Swiss-Prot/TrEMBL protein accession numbers. The protein intensity values were log_2_ transformed, followed by normalization across the five batches using the batch removal function in the LIMMA package. The normalized expression data were further transformed using Z-score procedure. The expression data were averaged across both sexes for each strain. Any proteins from Y chromosome were excluded from the dataset.

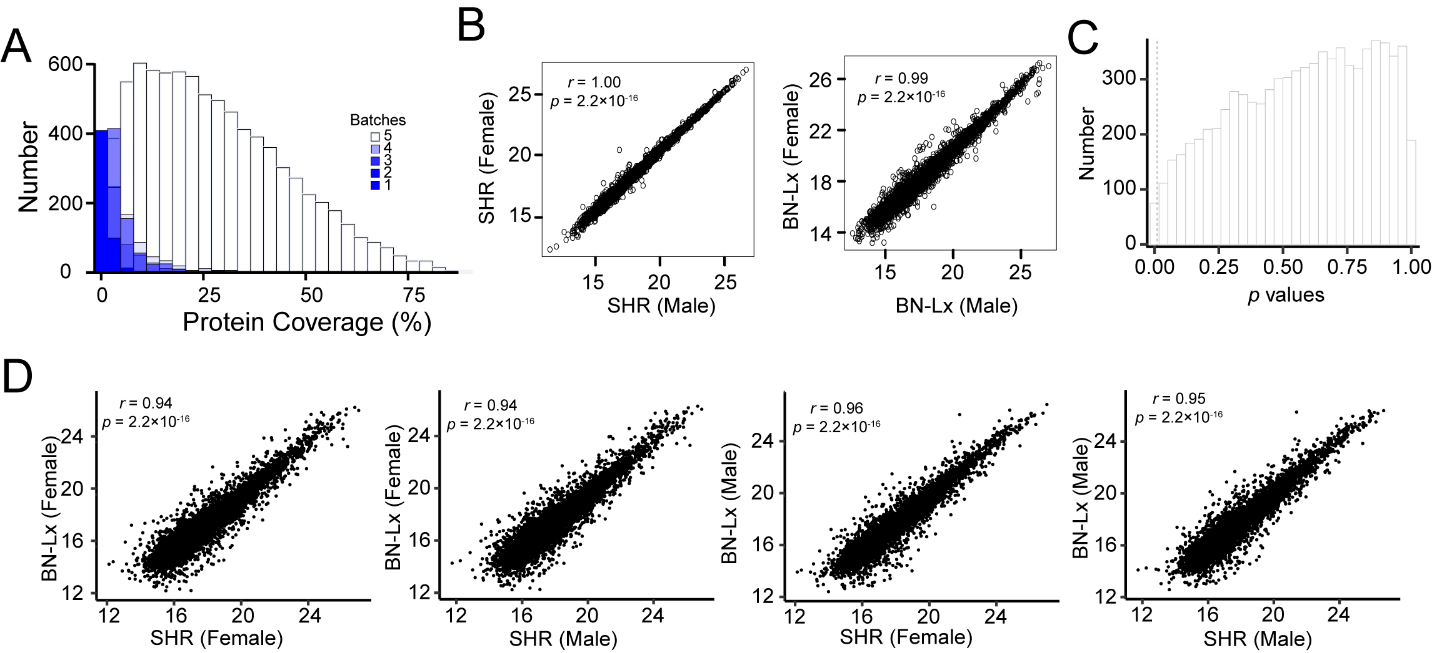

**Figure S2. Protein coverage and correlation analysis between two parental strains.**

(A) Histogram showing the coverage of quantified proteins across 29 batches of TMT experiments. The open bar represents the distribution of proteins quantified by all five TMT batches, while the blue gradient color shows the distribution of proteins detected in 1 to 4 TMT batches. Protein coverage is defined as the percentage of amino acids identified by the proteomics data. (B) Scatter plot displaying correlations between male and female of the SHR and BN-Lx strains. (C) Distribution of *p* values from the statistical tests comparing male and female samples for all proteins. (D) Scatter plots showing the correlations between the SHR and BN-Lx strains.

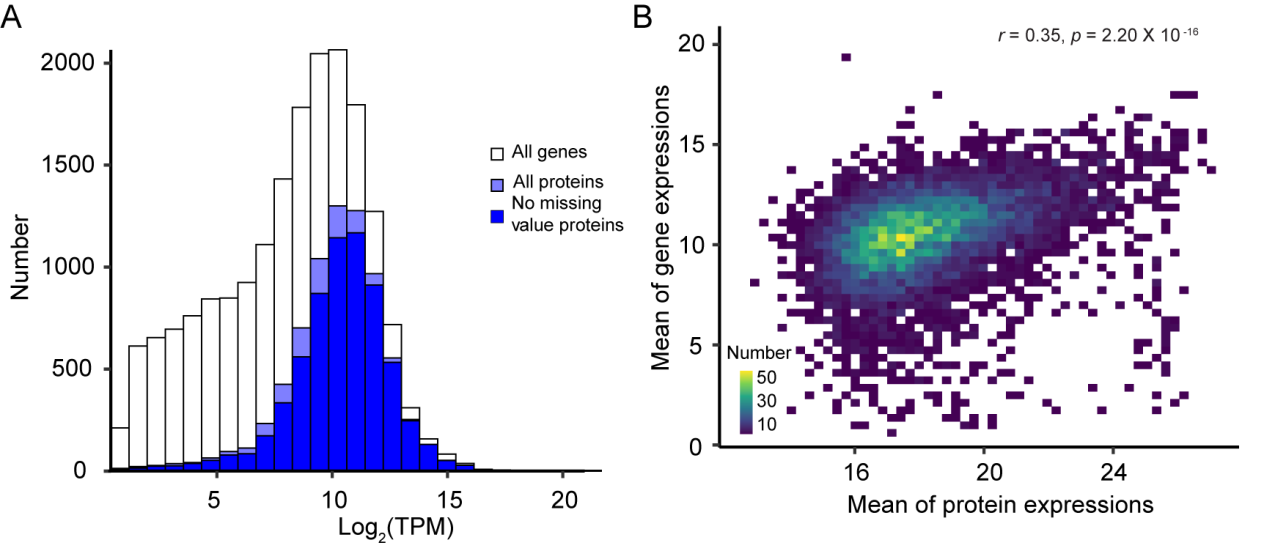

**Figure S3. Comparison between proteomic and transcriptomic data.**

(A) Histogram showing the coverage of proteomic data compared to RNA-seq data. The open bar represents the distribution of protein-coding genes detected by RNA-seq, the light blue bar indicates the distribution of protein-coding genes from proteomic data, and the navy bar indicates the distribution of protein-coding genes from no missing value proteomic data. (B) Scatter plot showing a comparison of gene expression levels and protein abundance. The expression is defined as the average expression across all samples.

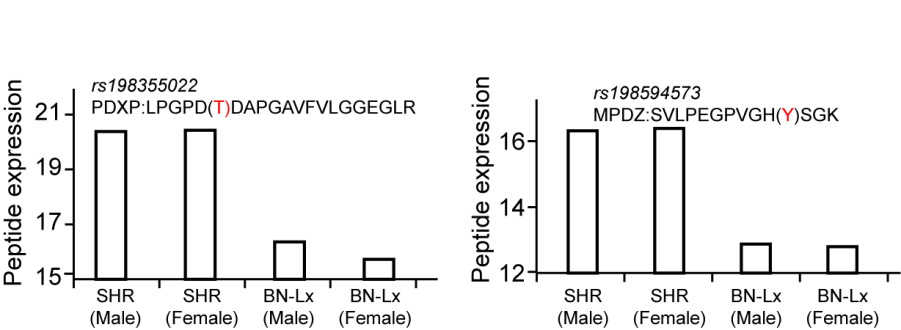

**Figure S4.** **Expression levels of variant peptides in SHR and BN-Lx strains.**

The amino acid highlighted in red within the parenthesis represents the reference amino acid encoded by the allele from the reference BN-Lx genome.

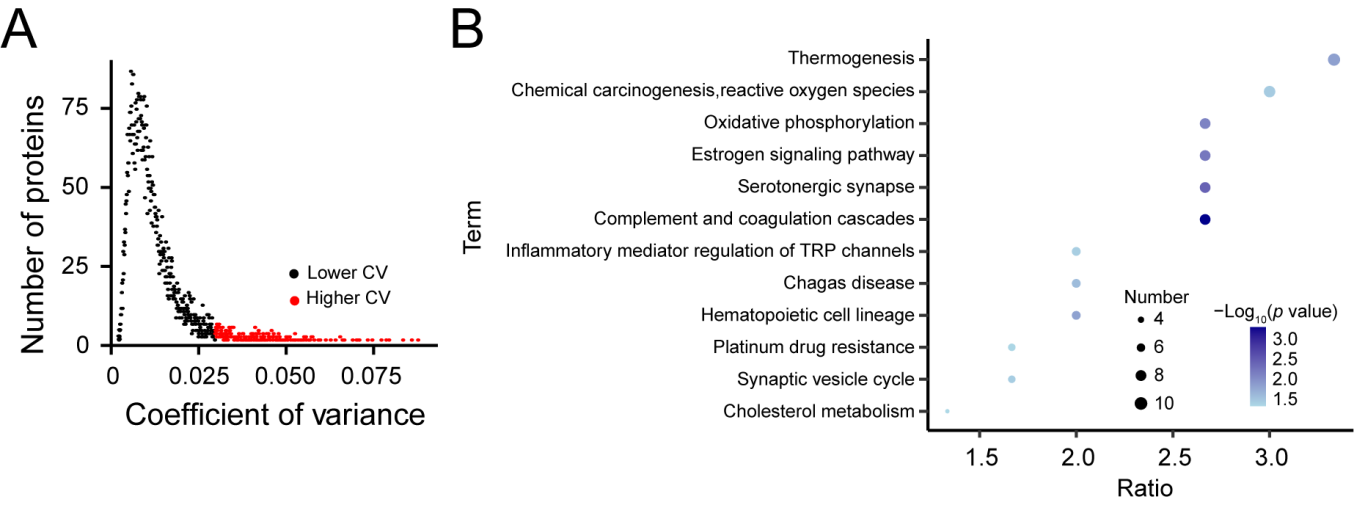

**Figure S5. Analysis of highly variable proteins across all 29 stains**.

(A) Distribution of coefficient of variation (CV) for all proteins across all samples. (B) Enriched KEGG pathways in the highly variable proteins.

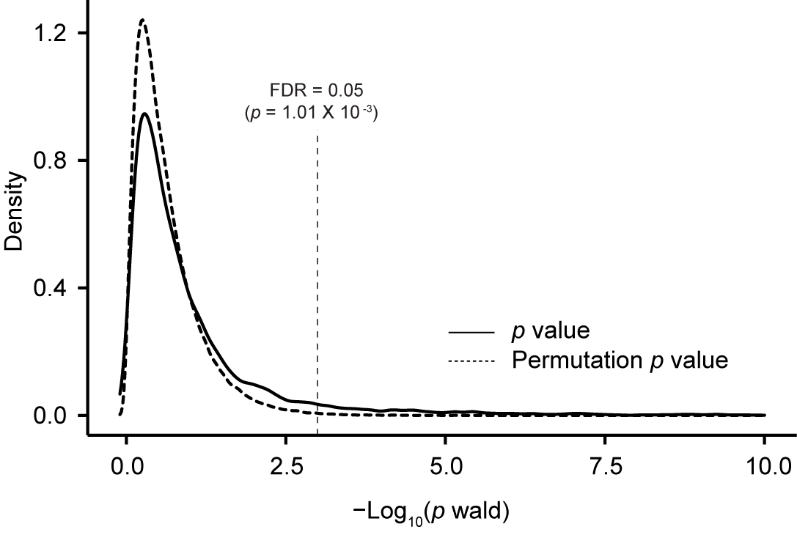

**Figure S6. Determination of the *p*-value cutoff.**

The *p* value cutoff is determined using the distributions of true *p* values and permutated *p* values.
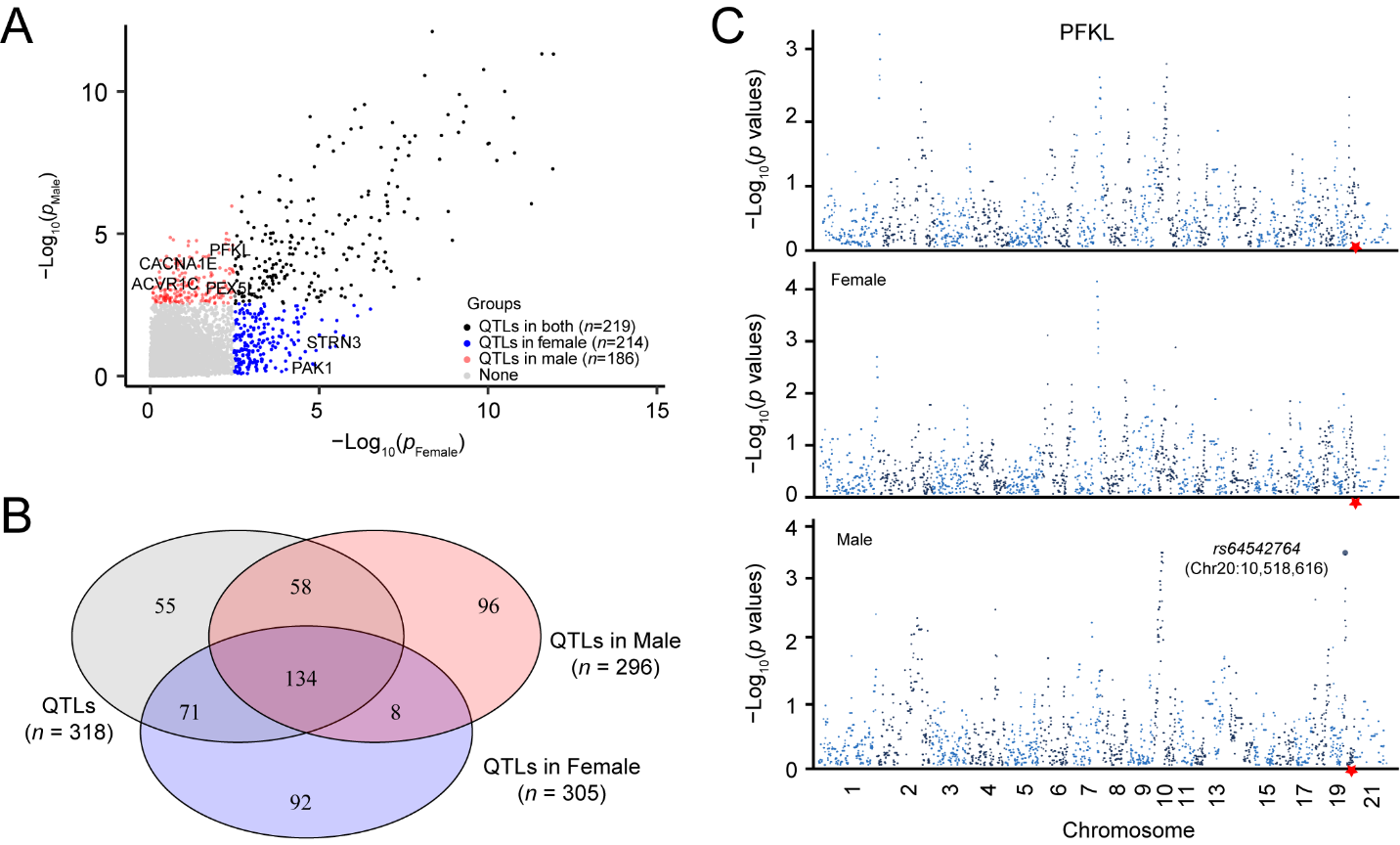

**Figure S7. Sex-specific genetic regulation of the brain proteome.**

(A) Scatter plot illustrating the significance of QTLs identified in both sexes, with QTLs specific to females (red), males (blue), and those found in both (black). (B) Venn diagram showing the overlap and distinct QTLs identified in male and female linkage analyses, as well as those common to both. (C) Manhattan plots display the genome-wide p-values for associations between SNPs and PFKL protein expression, separated by sex. The top *cis*-pQTL (*rs64542764*) for PFKL protein expression in males is indicated by a red diamond.

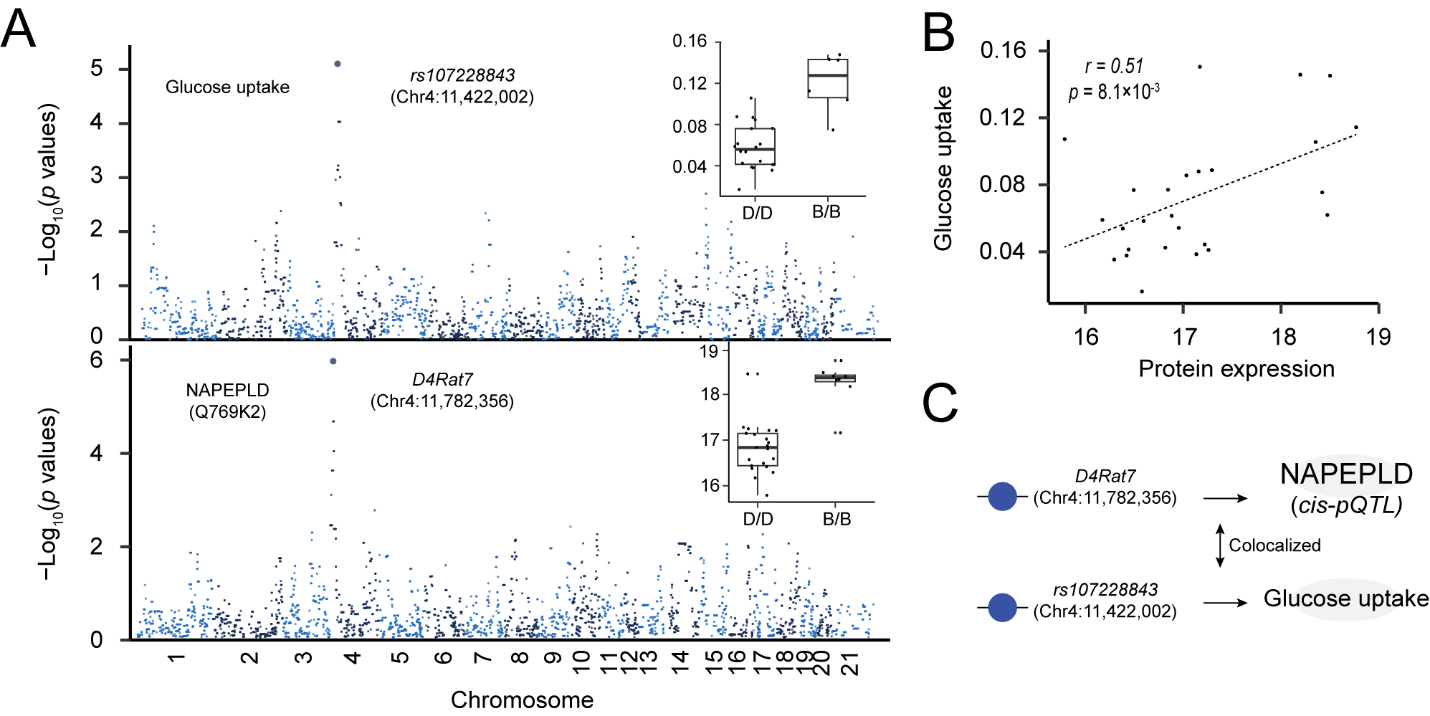

**Figure S8. Co-localization of *cis*-QTLs and phenotypic traits.**

(A) Manhattan plot showing a colocalized QTL associated with a phenotypic trait, glucose uptake, and associated with NAPEPLD protein expression. (B) Correlation of NAPEPLD expression level and glucose uptake. (C) Schematic representation illustrating the pleiotropic effect of the genotype on both protein expression and glucose uptake levels.

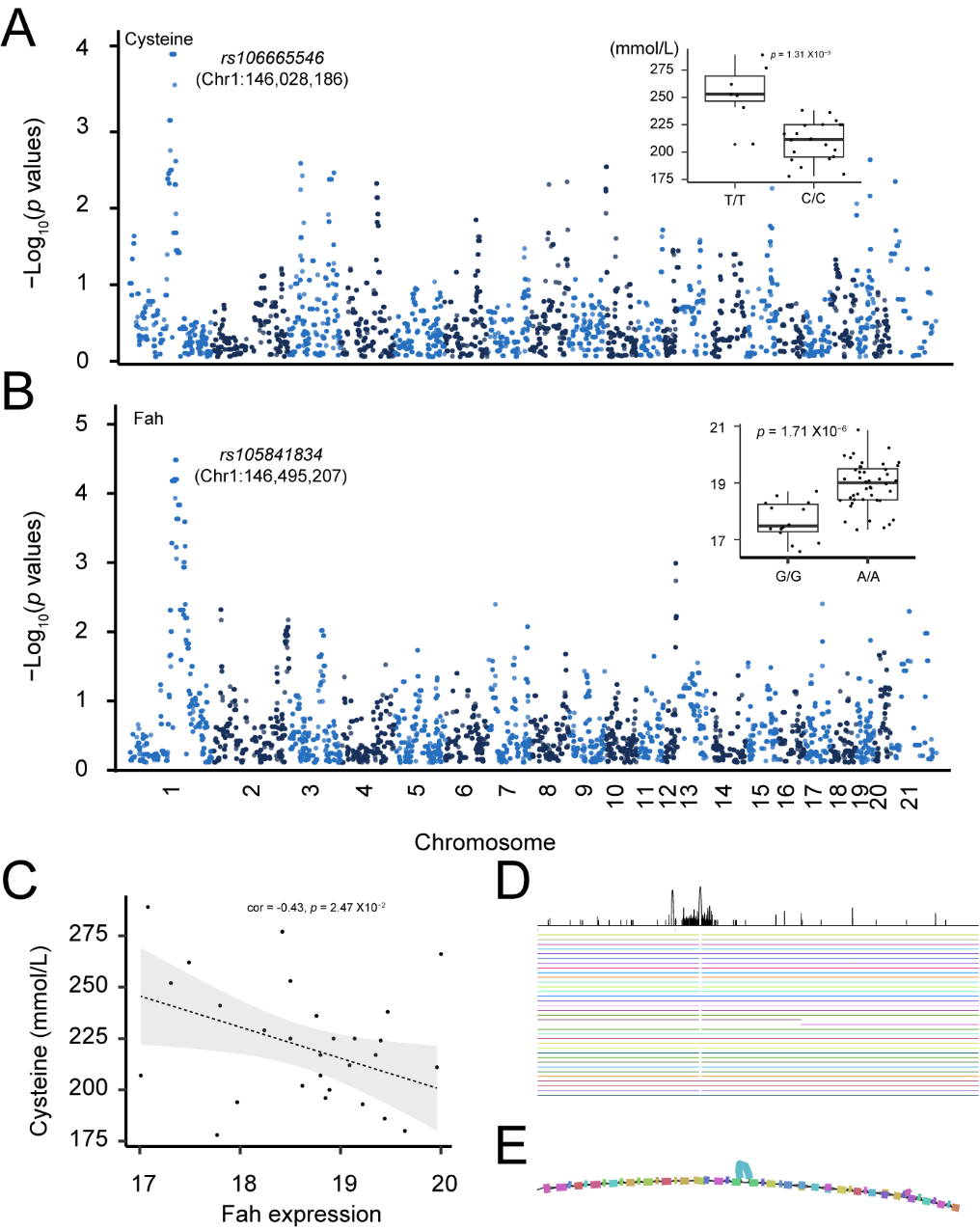

**Figure S9. Pangenome analysis revealing genetic variation among mapping strains.**

(A) Manhattan plot depicting genome-wide association of the locus *rs106665546* associated with blood cysteine levels, box plots showing the distribution of trait values by genotype. (B) Manhattan plot depicting genome-wide association of the locus *rs105841834* associated with Fah protein expression, box plots showing the distribution of trait values by genotype. (C) Scatter plot correlating Fah gene expression with blood cysteine levels. (D) Genome browser view displaying genetic variants within the Fah gene, with an evident large insertion highlighted in SHR and some HXB strains. (E) Schematic arc plot visualizing the structural variation of Fah genes across SHR and HXB/BXH strains.

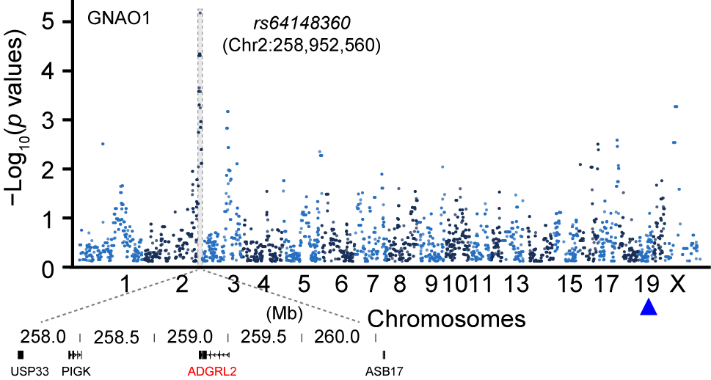

**Figure S10.** Manhattan plot showing a *trans*-pQTL (i.e., *rs64148360*) is associated with GNAO1 protein expression.

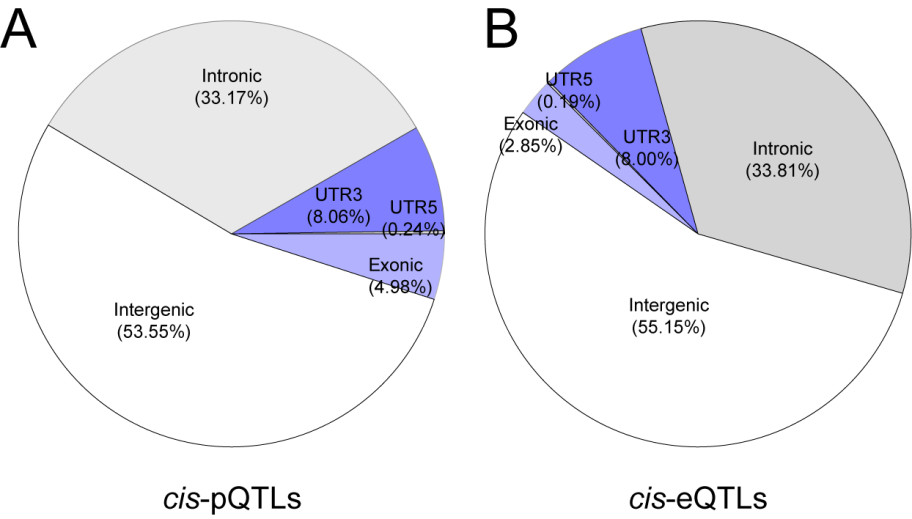

**Figure S11. Proportions of QTLs in different genomic regions.**

The pie chart is divided into five slices, each corresponding to a different genomic region: 5’ UTR, 3’ UTR, exon, intron, and intergenic. The size of each slice reflects the percentage of *cis*-pQTLs found in that region.
